# Mutant Huntingtin impairs neurodevelopment in human brain organoids through CHCHD2-mediated neurometabolic failure

**DOI:** 10.1101/2023.06.03.543551

**Authors:** Pawel Lisowski, Selene Lickfett, Agnieszka Rybak-Wolf, Stephanie Le, Werner Dykstra, Barbara Mlody, Carmen Menacho, Philipp Roth, Yasmin Richter, Linda A. M. Kulka, Petar Glazar, Haijia Wu, David Meierhofer, Ivano Legnini, Max Otto, Duncan Miller, Nancy Neuendorf, Tobias Hahn, Narasimha Swamy Telugu, Annett Böddrich, Ertan Mayatepek, Sebastian Diecke, Heidi Olzscha, Janine Kirstein, Ralf Kühn, Sidney Cambridge, Nikolaus Rajewsky, Erich E. Wanker, Josef Priller, Jakob J. Metzger, Alessandro Prigione

**Affiliations:** Max Delbrueck Center for Molecular Medicine (MDC), Berlin, Germany; Berlin Institute for Medical Systems Biology (BIMSB), Berlin, Germany; Neuropsychiatry, Charité - Universitätsmedizin Berlin, Germany; Institute of Genetics, Polish Academy of Sciences, Magdalenka, Poland; Department of General Pediatrics, Neonatology and Pediatric Cardiology, Medical Faculty, Heinrich-Heine-University, Duesseldorf, Germany; Cell Biology, University Bremen, Germany; Institute of Physiological Chemistry, Martin-Luther-University, Halle-Wittenberg, Germany; Max Planck Institute for Molecular Genetics, Berlin, Germany; Institute of Molecular Medicine, Medical School Hamburg, Germany; Berlin Institute of Health (BIH), Berlin, Germany; Leibniz Institute on Aging – Fritz-Lipmann Institute, Jena, Germany; Dr. Senckenbergische Anatomie, Goethe University Frankfurt, Germany; German Center for Neurodegenerative Diseases (DZNE), Berlin, Germany; Department of Psychiatry and Psychotherapy; School of Medicine and Health, Technical University Munich, Germany; University of Edinburgh and UK DRI, Edinburgh, UK

## Abstract

Expansion of the glutamine tract (poly-Q) in the protein Huntingtin (HTT) causes the neurodegenerative disorder Huntington’s disease (HD). Emerging evidence suggests that mutant HTT (mHTT) disrupts brain development. To gain mechanistic insights into the neurodevelopmental impact of human mHTT, we engineered induced pluripotent stem cells to introduce a biallelic or monoallelic mutant 70Q expansion or to remove the poly-Q tract of HTT. 70Q introduction caused aberrant development of cerebral organoids with loss of neural progenitor organization. The early neurodevelopmental signature of mHTT highlighted the dysregulation of the protein coiled-coil-helix-coiled-coil-helix domain containing 2 (CHCHD2), a transcription factor involved in mitochondrial integrated stress response. CHCHD2 repression was associated with abnormal mitochondrial morpho-dynamics and elevated resting energy expenditure. Elimination of the poly-Q tract of HTT normalized CHCHD2 expression and mitochondrial defects. Hence, mHTT-mediated disruption of human neurodevelopment is paralleled by aberrant neurometabolic programming mediated by dysregulation of CHCHD2, which could then serve as an early intervention target for HD.

## Introduction

Huntington’s disease (HD) is a rare neurodegenerative disorder caused by inherited defects in the gene Huntingtin (*HTT*) encoding for the protein HTT. The mutant *HTT* (mHTT) gene exhibits an abnormal (>35) CAG expansion resulting in elongated repeats of glutamine (Q) in the poly-Q tract of the protein ^1^. HD is currently incurable and the mechanisms underlying the neurodegenerative process are not fully understood.

An increasing amount of evidence points towards impaired brain development in HD. In fact, although wild-type (WT) HTT is present in several tissues, it is expressed at highest level in the brain, even before the completion of neuronal maturation ^2^. In mice, targeted HTT disruption is embryonic lethal ^3, 4^, brain-specific HTT inactivation leads to progressive neuronal defects ^5^, and extensive HTT reduction fails to support normal brain development ^6^. WT HTT may thus exert a physiological role in neurodevelopment, including axonal transport ^7^, synapse development ^8^, neural rosette formation ^9^, neuronal migration ^10^, as well as regulation of neural progenitor cells (NPCs) and neurogenesis ^11^.

In accordance with a developmental component in HD, mHTT with poly-Q tract longer than 60 causes severe juvenile forms of HD, which recapitulate features associated with neurodevelopmental disorders ^12^. Even individuals with non-juvenile forms of HD can develop signs of impaired brain growth during childhood before the occurrence of clinical manifestation ^13–15^. In mice, mHTT disrupts the division of cortical progenitors ^16^, and temporally-limited expression of mHTT in early life is sufficient to recapitulate the disease phenotypes ^17^. Importantly, the analysis of human fetuses carrying mHTT demonstrated similar neurodevelopmental defects, with diminished numbers of proliferating NPCs and higher number of NPCs that prematurely enter neuronal lineage specification ^18^.

An effective model system for dissecting the neurodevelopmental aspects of HD is represented by human induced pluripotent stem cells (iPSCs). The transcriptional signature of neurons differentiated from iPSC models of HD pointed toward a dysregulation of neurodevelopmental genes ^19–21^. Cerebral organoids carrying mHTT exhibited aberrant organization and specification of NPCs, with immature ventricular zones ^22^. The CAG repeat length could thus regulate the balance between NPC expansion and differentiation in cerebral organoids ^23^. Micro-patterned neuruloids consisting of NPCs, neural crest, sensory placode and epidermis showed disrupted self-organization in the presence of mHTT, indicating faulty neurodevelopment ^24^.

In order to gain mechanistic insights into how mHTT impacts human brain development, we generated brain organoids from engineered isogenic iPSC lines in which we introduced mHTT (70Q) in one or both *HTT* alleles, or we eliminated the poly-Q stretch completely from the HTT gene. We found that biallelic 70Q introduction disrupts the development of brain organoids (unguided cerebral organoids and region-specific cortical organoids and midbrain organoids) causing defective NPC organization despite the presence of neuronal generation. To identify mechanisms regulating early developmental defects caused by mHTT, we focused on shared signatures across undifferentiated iPSCs and neural committed cells. We identified the mitochondrial protein coiled-coil-helix-coiled-coil-helix domain containing 2 (CHCHD2) as a top dysregulated factor. In accordance to the known function of CHCHD2 in mitochondrial integrated stress response (mISR) ^25^, neural cells carrying mHTT developed a mISR signature with disruption of mitochondrial dynamics and bioenergetic alterations characterized by increased energy expenditures at rest, a condition known as hypermetabolism. iPSCs and NPCs responded to mitochondrial dysfunctions by compensatory upregulation of glycolysis, but pure neuronal cultures from HD patient-derived iPSCs reduced both mitochondrial and glycolytic metabolism. In frame elimination of the poly- Q tract in HTT reverted the defects in CHCHD2 expression and mitochondrial morpho- dynamics. Hence, we were able to link mHTT to altered mitochondrial function and metabolism through the dysregulation of CHCHD2, which could lead to the observed developmental defects.

## Results

### mHTT impairs early NPC organization in engineered human cerebral organoids

We engineered iPSCs with a healthy genomic background (WT/WT) using CRISPR/Cas9 ^26, 27^. The edited region within the *HTT* gene entailed both the CAG repeat stretch (poly-Q in the protein) and the CCG repeat stretch (poly-P in the protein) **(Fig. 1a, Supplementary Fig. 1a)**. As template, we used an expanded tract composed of 70 CAG/CAA repeats mimicking mutations observed in patients. The resulted engineered iPSC lines were: i) WT/70Q carrying the expanded CAG tract only on one allele, ii) 70Q/70Q carrying the expanded CAG tract on both alleles, iii) 0Q/0Q with an in-frame deletion that eliminated the CAG and CCG stretches **(Fig. 1a-b)**. We confirmed the successful editing by sequencing, PCR, and immunoblotting **(Fig. 1c, Supplementary Fig. 1b-f)**. The set of isogenic iPSC lines exhibited a normal karyotype and displayed no modification of top computationally predicted off-target sites **(Supplementary Fig. 1g-h)**.

**Fig. 1.**
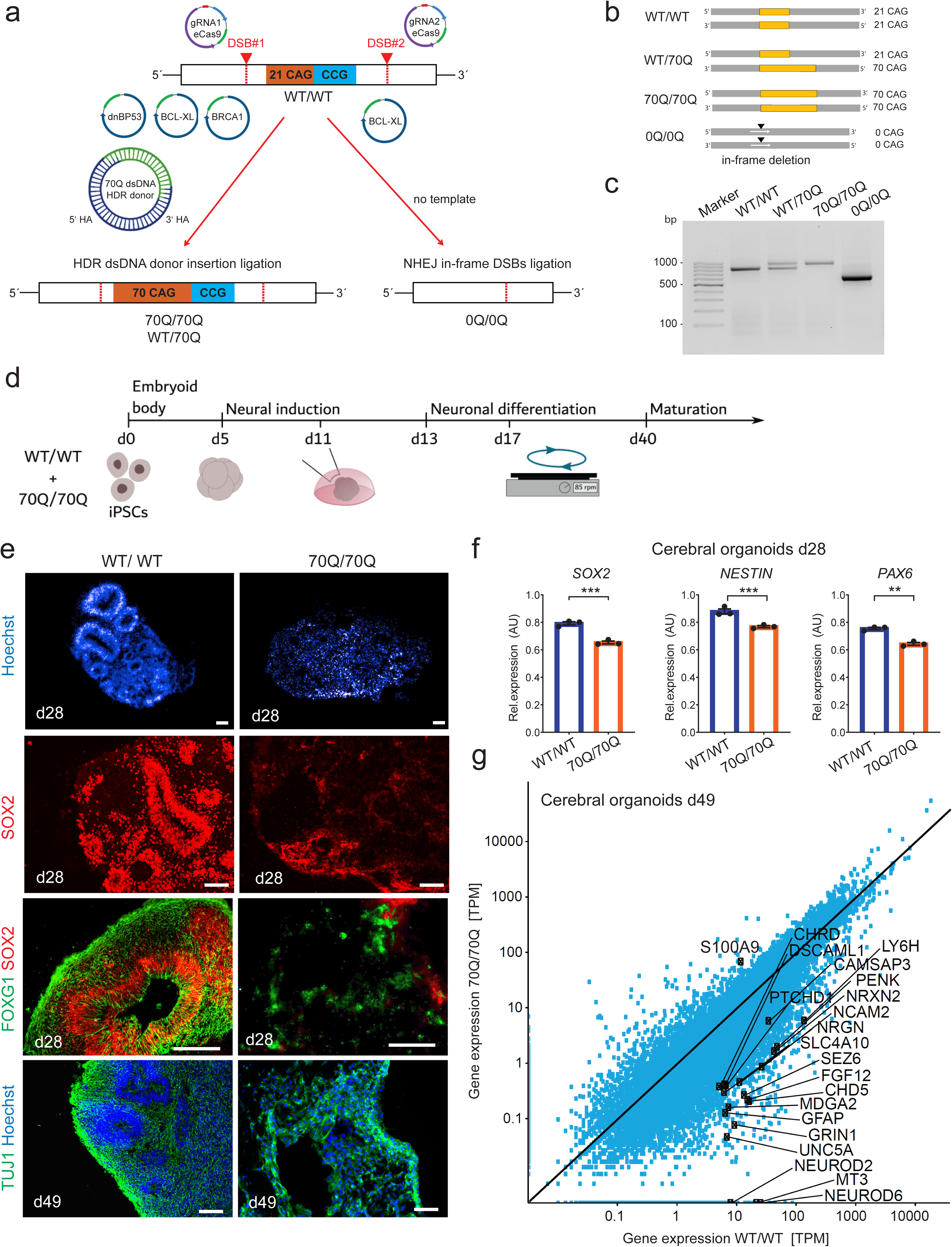
Engineered iPSCs carrying mHTT give rise to neurodevelopmentally impaired cerebral organoids. **a** Genome editing approach in control iPSCs (WT/WT) using two double-strand breaks (DSB) sites to modify the HTT genomic region encompassing the CAG/CAA and CCG repeats stretches. To generate iPSCs carrying elongated CAG in one allele (70Q/WT) or in both alleles (70Q/70Q), we promoted homology direct repair (HDR) (with plasmids BCL-XL and BRCA1), inhibited non-homologous end joining (NHEJ) (with plasmid dnBP53), and provided a HDR donor dsDNA plasmid carrying 70Q repeats and homology arms. We harnessed NHEJ to obtain iPSCs with in-frame deletion of the CAG/CCG region (0Q/0Q). **b** Overview of the engineered isogenic iPSC lines. **c** PCR analysis of HTT in the isogenic iPSC lines. **d** Schematics of the protocol to generate unguided cerebral organoids from isogenic iPSC lines WT/WT and 70Q/70Q. **e** Immunostaining in cerebral organoids at day 28 and 49 showing defective cytoarchitecture and neural progenitor cell (NPC) organization in 70/70Q. Scale bar: 100 µm. **f** qPCR analysis of NPC markers in cerebral organoids at day 28. Mean +/- s.e.m.; n=3 independent biological replicates (dots) per line; ***p<0.001 WT/WT vs. 70Q/70Q; unpaired two-tailed t test. Four organoids were pooled for each individual RNA isolation. **g** Gene expression analysis of cerebral organoids at day 49 highlighting the genes belonging to the GO term “nervous system development” (GO:0007399).

In order to assess the impact of mHTT on human brain development, we focused on the line 70Q/70Q as an example of strong mHTT pathology. While heterozygous cell lines with expanded poly-Q tract have previously been used for the analysis of mHTT effects, we employed here the homozygous 70Q/70Q line to identify the most severe consequences of mHTT on human neurodevelopment without compensatory effects of WT HTT. To our knowledge, a homozygous model of mHTT has not yet been studied using iPSCs. Using this model (70Q/70Q) and its isogenic control (WT/WT), we applied an unguided cerebral organoid differentiation protocol **(Fig. 1d)**. Cerebral organoids appeared strongly affected by mHTT. The overall cellular organization was disrupted in 70Q/70Q cerebral organoids, with a striking lack of ventricular zone-like neurogenic zones **(Fig. 1e)**. NPCs were particularly impaired, as demonstrated by the reduced presence and spatial organization of cells positive for NPC markers SOX2 and FOXG1 **(Fig. 1e)**. In accordance, transcriptional analysis of cerebral organoids at different time points of development showed the reduction of progenitor markers **(Fig. 1f)**. Additionally, the great majority of gene associated with the GO term “nervous system development” were downregulated in mTT-expressing organoids **(Fig. 1g)**. Despite these dramatic alterations in the progenitor population, we could still detect neuronal cells, as indicated by the markers TUJ1, MAP2, and SMI32 **(Fig. 1e, Supplementary Fig. 2a-b)**. These findings are in agreement with previous data of mHTT-carrying human fetuses reporting the loss of proliferating NPCs and premature neuronal differentiation ^18^.

To dissect the impact of developmental alterations in different brain regions, we generated guided region-specific brain organoids **(Fig. 2a)**. We focused on cortical organoids and midbrain organoids. In fact, according to neuroanatomical reports, neurodegeneration in HD individuals may not be limited to striatum and may affect also brainstem and neocortex regions ^28^. Under physiological conditions, cortical organoids typically reach a much larger volume over time than midbrain organoids, which instead remain relatively small and are kept under static growth ^29^. We found that the presence of mHTT strongly diminished the size development of both cortical and midbrain organoids **(Fig. 2b-c)**. In both region-specific organoids, the overall size appeared altered already at relatively early stages, even before 10 days of differentiation **(Fig. 2b-c)**. We next checked for presence of toxic HTT aggregates during organoid development. We could not find significant presence of large-size HTT protein conformations after 70 days of cultures **(Supplementary Fig. 2c-d)**. Collectively, these data suggest that mHTT causes neurodevelopmental defects that occurs early on during development, and impact neural progenitors well before the accumulation of potentially toxic protein aggregates.

**Fig. 2.**
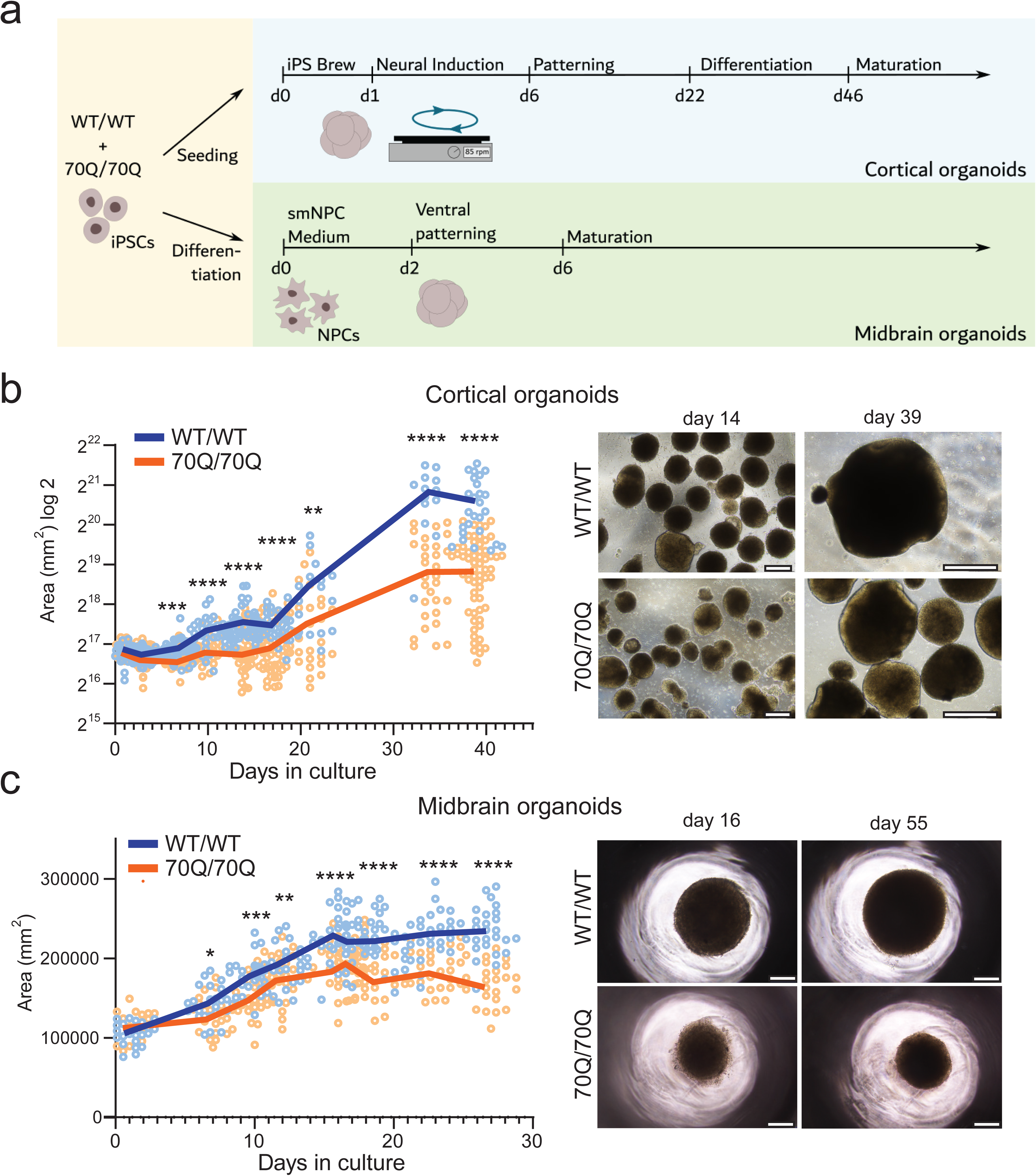
mHTT compromises the development of region-specific brain organoids. **a** Schematics of the protocol to generate guided region-specific brain organoids from isogenic iPSC lines WT/WT and 70Q/70Q. For cortical organoids, we started the differentiation from iPSCs, for midbrain organoids we started from NPCs. **b** Left: growth curve of cortical organoids. Dots represent individual organoids over two independent experiments. We compared organoid size at defined time points (day 1, 2, 7, 10, 14, 17, 21, 34, 39): ***p<0.001, ****p<0.0001 WT/WT vs. 70Q/70Q; two-tailed Mann-Whitney U test. Right: examples of cortical organoids at bright field microscope. Scale bar: 400 µm. **c** Left: growth curve of midbrain organoids. Dots represent individual organoids over two independent experiments. We compared organoid size at defined time points (day 1, 7, 10, 12, 16, 17, 19, 23, 27): ***p<0.001, ****p<0.0001 WT/WT vs. 70Q/70Q; two-tailed Mann-Whitney U test. Right: examples of cortical organoids at bright field microscope. Scale bar: 400 µm.

### CHCHD2 is a top dysregulated gene within the early neurodevelopmental signature of mHTT

We next aimed to transcriptionally dissect the mechanisms underlying the early neurodevelopment defects caused by mHTT since polyQ tract motifs are present in transcription factors and act as transcriptional regulating domains through mediation of binding between transcription factors and transcriptional regulators {Mohan, 2014 #1042}. For clear transcriptional dissection of mHTT we carried out total RNA sequencing of 70Q/70Q and WT/WT at different time points of neurodevelopment: iPSCs, NPCs, and cerebral organoids at day 28 and day 49, which are time points at which neural progenitors are already being formed **(Fig. 3a, Supplementary Data 1)**. We focused on genes that were dysregulated early and continuously throughout these differentiation time points. We identified a common expression signature composed of 47 genes that were either down- or upregulated in 70Q/70Q compared to WT/WT **(Fig. 3a, Supplementary Fig. 3a-b, Supplementary Data 2).** Within this signature, the most downregulated genes in mHTT-expressing cells was coiled-coil-helix-coiled-coil-helix domain containing 2 (CHCHD2) **(Fig. 3b)**. Other downregulated factors included genes involved in development (GBX2, PAX7, PAX8, LHX1, LMX1B, BARX1), neuronal development (POUF33, POUF3F2, TAF9B, EN2), and glutamate metabolism (SLC1A2, GAD1, GAD2) **(Fig. 3b, Supplementary Fig. 3b)**. Upregulated factors in mHTT-expressing cells were genes associated with detoxification (GSTM1, IAH1), protein translation (EIF1AY, KDM5D), and protein degradation (USP9Y). In addition to this common signature, genes involved in nervous system development were consistently downregulated in NPCs and cerebral organoids carrying mHTT **(Supplementary Fig. 3d)**.

**Fig. 3.**
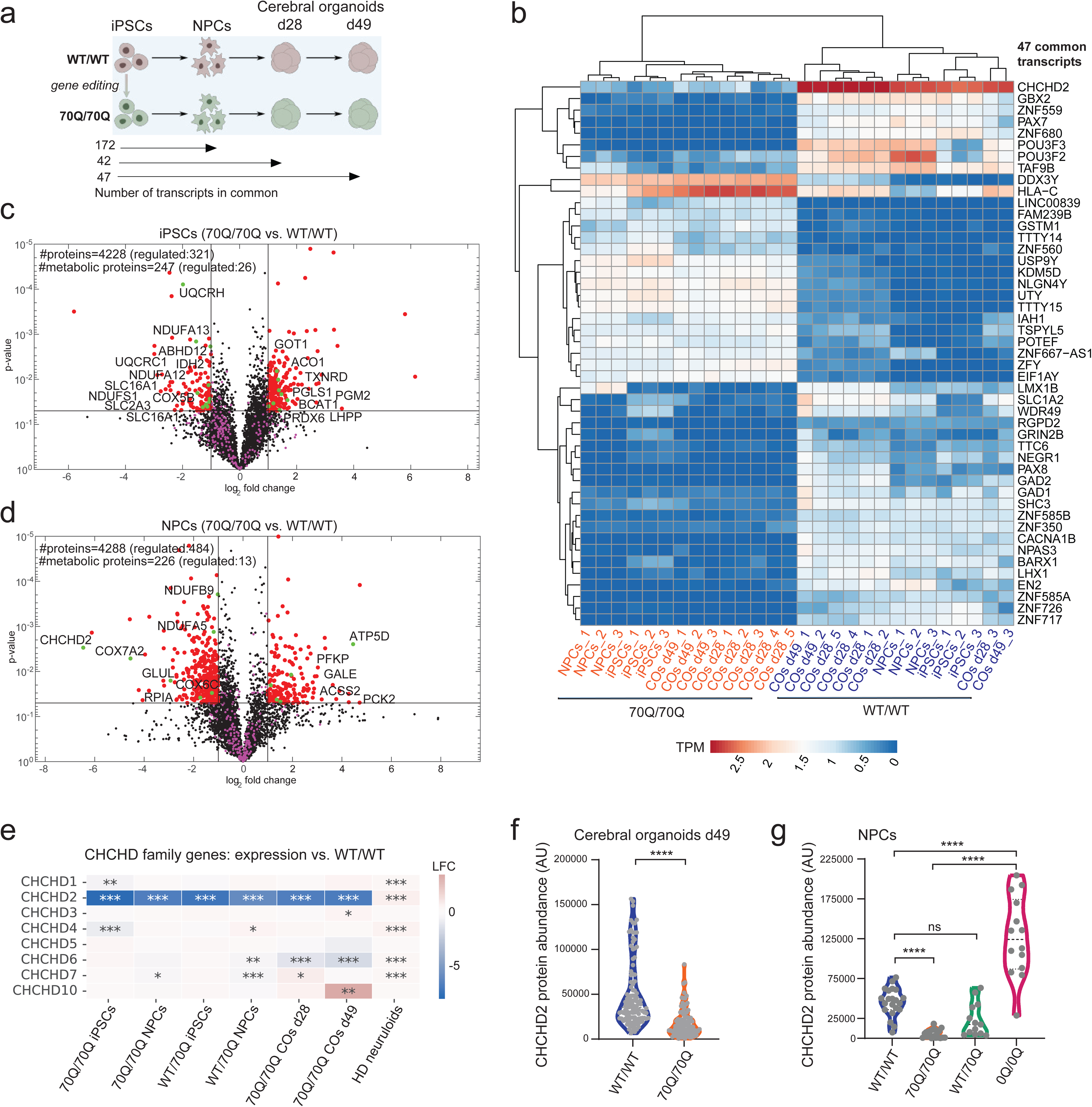
Omics analyses of mHTT-expressing cells across neurodevelopmental stages highlight the dysregulation of CHCHD2. **a** Schematics of RNA sequencing experimental set up for isogenic iPSC lines WT/WT and 70Q/70Q. Below are reported the number of transcripts that are uniquely in common between the different neurodevelopmental stages (see Supplementary Data 1-2). **b** Heatmap showing the 47 transcripts that are uniquely in common across all neurodevelopmental stages: iPSCs, NPCs, cerebral organoids (COs) at day 23 and COs at day 49. TPM: transcripts per million. **c-d** Volcano plot of proteomic datasets in iPSCs and NPCs showing statistical significance (p-value) versus magnitude of change (log_2_ fold change) (see Supplementary Data 3-5). Red dots indicate the significantly differently regulated proteins (right quadrant: upregulated in 70Q/70Q; left quadrant: downregulated in 70Q/70Q). Metabolic proteins were later used for the proteomic-driven functional metabolic analysis (see Supplementary Data 5). Green dots highlights those regulated proteins that are explicitly named. **e** Heatmap of log-fold change (LFC) comparisons of differential gene expression of genes belonging to the CHCHD family. Data were obtained from bulk RNA sequencing from iPSCs, NPCs, and COs derived from 70Q/70Q and WT/70Q compared to the respective cell type from WT/WT. HD neuruloids were previously published ^24^, and were compared to WT neuruloids. **f-g** Quantifications of protein abundance of CHCHD2 based on immunostaining performed in cerebral organoids and NPCs. The amount of positive CHCHD2 signal per image was normalized to the Hoechst signal. Dots represent individual images collected over two independent experiments. ****p<0.0001, ns: not significant; two-tailed Mann-Whitney U test.

We then performed global proteomics of iPSCs and NPCs to determine whether any of the observed transcriptional changes induced by mHTT were recapitulated at the protein level. We identified metabolism-related proteins dysregulated in both iPSCs (46 metabolic proteins out 321) and NPCs (13 out of 484) **(Fig. 3c-f, Supplementary Data 3-4)**. Among these proteins, CHCHD2 was significantly downregulated in 70Q/70Q NPCs **(Fig. 3d)**.

Since both transcriptomics and proteomics identified CHCHD2 as an important dysregulated target upon 70Q introduction, we inspected the transcription level of the whole family of coiled-coil-helix-coiled-coil-helix (CHCH) domain-containing proteins. These are proteins imported into the mitochondrion that have been suggested to play a role in neurodegenerative diseases ^30, 31^. Although other CHCH family members were dysregulated by mHTT, CHCHD2 appeared most significantly affected in both iPSCs and NPCs carrying the elongated CAG repeat in both alleles (70Q/70Q) or in only one allele (WT/70Q) **(Fig. 3e)**. Using previously published datasets of neuruloids carrying mHTT ^24^, we found that also neuruloids showed dysregulation of CHCHD2 **(Fig. 3e).**

In order to validate the defective expression of CHCHD2 in cells carrying mHTT, we performed immunostaining for CHCHD2 in cerebral organoids and NPCs and applied an automated pipeline to quantify the expression of CHCHD2. Using this approach, we confirmed a decreased level of CHCHD2 protein in cerebral organoids and NPCs carrying mHTT (70Q/70Q) compared to WT/WT **(Fig. 3f-g)**. NPCs carrying mHTT in only one allele (WT/70Q) also showed an overall reduced abundance of CHCHD2, but this difference did not reach statistical significance **(Fig. 3g)**. Remarkably, the elimination of both poly-Q and poly-P tracts (0Q/0Q) resulted in a significant increase of CHCHD2 protein expression in NPCs **(Fig. 3g)**. Hence, mHTT leads to downregulation of CHCHD2 that can be reverted upon eliminating the CAG/CCG region. These findings point towards a potential role for CHCHD2 dysregulation in HD pathogenesis.

### CHCHD2 dysregulation by mHTT is associated with a mISR signature and defective mitochondrial morpho-dynamics

We next investigated the downstream consequences associated with the dysregulation of CHCHD2. CHCHD2 is a mitochondrially imported protein that has also been reported to act as a transcription factor and translocates to the nucleus ^32^. In fact, CHCHD2 appeared in the list of nucleic acid binding genes **(Supplementary Fig. 3c)**. Using semi-automated quantification of immunostained images, we confirmed that CHCHD2 co-localization with Hoechst was significantly diminished in both NPCs and cerebral organoids carrying mHTT 70Q/70Q **(Fig. 4a-b)**.

**Fig. 4.**
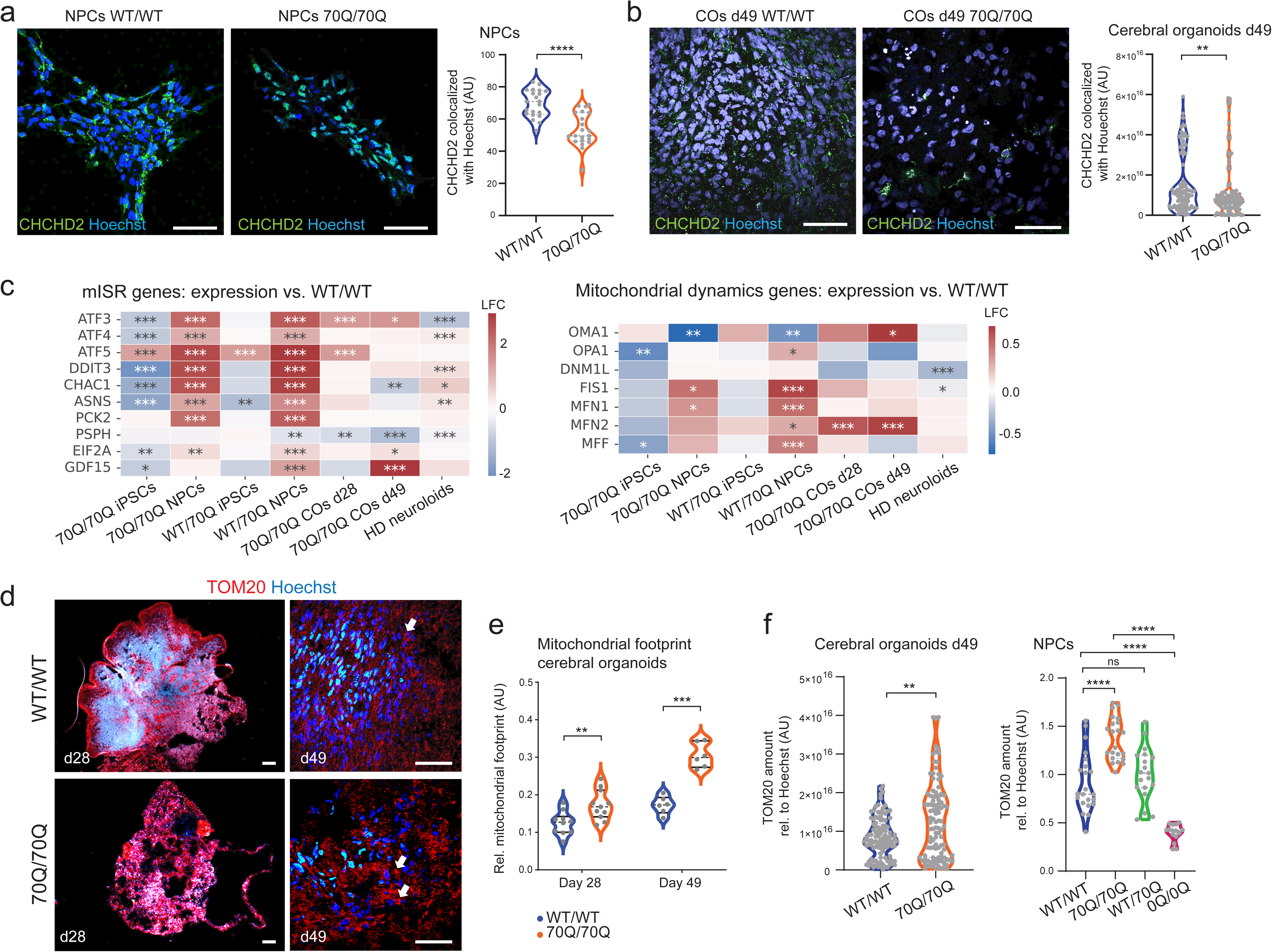
Mitochondrial integrated stress response (mISR) and altered mitochondrial network in mHTT-carrying neural cells where CHCHD2 is dysregulated. **a-b** Representative immunostaining images and related quantifications of CHCHD2 protein colocalization with nuclear staining Hoechst in NPCs and cerebral organoids. Dots represent individual images collected over two independent experiments. **p<0.01, ****p<0.0001; two-tailed Mann-Whitney U test. Scale bar: 100 µm. **c** Heatmap of log-fold change (LFC) comparisons of genes involved in mitochondrial integrated stress response (mISR) and mitochondrial dynamics based on bulk RNA sequencing from iPSCs, NPCs, and cerebral organoids (COs) derived from 70Q/70Q and WT/70Q compared to the respective cell type from WT/WT. HD neuruloids ^24^ were compared to WT neuruloids. **d-e** Representative immunostaining images and related quantification of mitochondrial footprint (comprising both small and large mitochondrial structures) based on TOM20 signal in cerebral organoids from 70Q/70Q and WT/WT. Dots represent individual images collected over two independent experiments. **p<0.01, ***p<0.001, unpaired two-tailed t test. Scale bar: 100 µm. **f** Quantifications of protein abundance of TOM20 based on immunostaining performed in cerebral organoids and NPCs. The amounts of positive TOM20 signal per image was normalized to Hoechst signal. Dots represent individual images collected over two independent experiments. **p<0.01, ****p<0.0001, ns: not significant; two-tailed Mann-Whitney U test.

CHCHD2 plays a role in mitochondrial integrated stress response (mISR) ^25^. Accordingly, genes associated with mISR were dysregulated upon 70Q introduction **(Fig. 4c)**. This was particularly evident for NPCs (70Q/70Q and WT/70Q), where mISR-related genes were significantly upregulated compared to WT/WT (e.g. ATF3, ATF4, DDIT3, CHAC1, PCK2). Altered mISR genes were present also in cerebral organoids at day 49 and in HD neuruloids, although to a lower extent. In conditions with impaired mitochondrial oxidative phosphorylation (OXPHOS), mISR is associated with dysregulation of mitochondrial quality control and mitochondrial dynamics, and cellular senescence ^33^. Indeed, genes related to mitochondrial quality control, mitochondrial biogenesis, and senescence were affected by mHTT, with particular upregulation of quality control genes in NPCs (70Q/70Q and WT/70Q) and cerebral organoids (70Q/70Q) (e.g. YMEIL1, LONP1, LONP2, HSPA1) **(Supplementary Fig. 4a)**. mHTT also altered the expression of genes regulating mitochondrial dynamics in mutant NPCs and cerebral organoids, with upregulation of genes related to mitochondrial fusion (e.g. FIS1, MNF1, MNF2) and mitochondrial fission (OMA1, MFF) **(Fig. 4c)**.

We next evaluated the mitochondrial network morphology in cerebral organoids based on the expression pattern of the mitochondrial outer membrane marker TOM20 **(Fig. 4d)** ^34^. In agreement with altered expression of genes involved in mitochondrial dynamics, mutant cerebral organoids exhibited elevated mitochondrial footprint, which was composed of both small mitochondrial structures and large mitochondrial structures **(Fig. 4e)**. This increase was more evident at day 49 compared to day 28, suggesting that defects in mitochondrial dynamics might become more severe over time. We also observed increased presence of hyper-fused mitochondrial structures in mutant organoids (**Fig. 4d,** white arrows). Using our semi-automated quantification of immunostaining images, we found higher relative levels of TOM20 protein in NPCs and cerebral organoids carrying 70Q/70Q **(Fig. 4f),** while this increase did not reach significance in NPCs with WT/70Q. Remarkably, elimination of the poly-Q/poly-P region was sufficient to repress the abnormal TOM20 increase **(Fig. 4f)**. Taken together, CHCHD2 dysregulation following 70Q introduction causes neural cells to develop a mISR signature and aberrant mitochondrial morpho-dynamics that can be reverted upon elimination of the CAG/CCG repeat region.

### mHTT causes hypermetabolism and aberrant metabolic programming

Following the identification of defects in mitochondrial morpho-dynamics caused by mHTT, we aimed to elucidate the associated functional metabolic consequences. Using the proteomics datasets, we carried out functional metabolic pathway analyses, as it was previously done to pinpoint a disruption in mitochondrial metabolism in cerebral organoids from autism spectrum disorders ^35^. This kinetic model comprises the major cellular metabolic pathways of energy metabolism in neural cells, including ion membrane transport and mitochondrial membrane potential **(Supplementary Data 5)**.

Both iPSCs and NPCs carrying mHTT showed higher glucose utilization at rest **(Fig. 5a)**. This feature of elevated resting energy expenditure (REE) is also known as hypermetabolism, a condition developing upon impaired OHXPHOS function in primary mitochondrial diseases that is associated with mISR activation ^33^. In mutant iPSCs, the maximal substrate utilization was reduced as was glucose uptake **(Supplementary Fig. 4b- c)**. mHTT also decreased the ATP/ADP ratio and maximal ATP production in iPSCs **(Supplementary Fig. 4b-c)**. In contrast, we did not observe differences in substrate utilization under augmented capacity or in glucose uptake in mHTT-expressing NPCs **(Supplementary Fig. 4b-c)**. 70Q introduction in both iPSCs and NPCs led to higher lactate production and diminished mitochondrial oxidation **(Fig. 5b, Fig. 5d)**. Accordingly, gene set enrichment analysis of global proteomics indicated that both mutant iPSCs and NPCs exhibited lower OXPHOS and higher glycolysis/gluconeogenesis **(Fig. 5c, Fig. 5e)**.

**Fig. 5.**
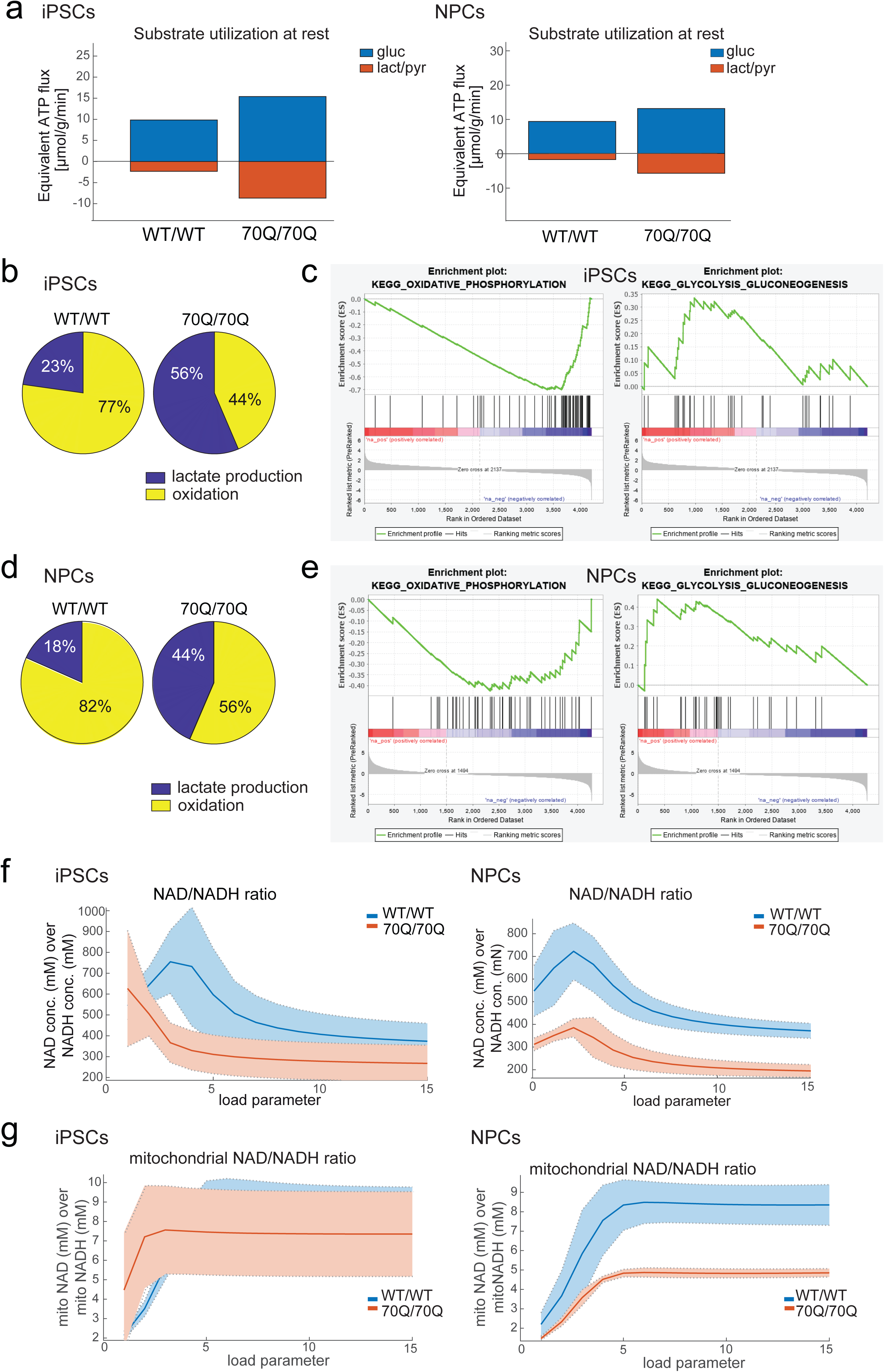
Neurometabolic defects induced by mHTT. **a** Proteomic-driven functional metabolic analysis (see Supplementary Data 5) depicting the substrate utilization at rest of iPSCs and NPCs from WT/WT and 70Q/70Q. Energetic capacities were evaluated by computing the changes of metabolic state elicited by an increase of the ATP consumption rate above the resting value. mHTT-carrying cells showed higher consumption of glucose (gluc) and higher production of lactate-pyruvate (lact/pyr). **b** Relative glucose utilization at resting energy demands in iPSCs based on functional metabolic analysis. **c** Gene Set Enrichment Analysis (GSEA) showing decreased oxidative phosphorylation and increased glycolysis/gluconeogenesis in 70Q/70Q iPSCs compared to WT/WT iPSCs. **d** Relative glucose utilization at resting energy demands in NPCs based on functional metabolic analysis. **e** GSEA showing decreased oxidative phosphorylation and increased glycolysis/gluconeogenesis in 70Q/70Q NPCs compared to WT/WT NPCs. **f-g** Metabolic state variables for NAD/NADH ratio and mitochondrial NAD/NADH ratio in iPSCs and NPCs in dependence of increasing energetic demands.

We next analyzed the ratio of NAD/NADH and mitochondrial NAD/NADH, which are important indicators of metabolic and redox homeostasis ^36^. Both ratios were clearly reduced in mutant NPCs compared to WT NPCs, indicating the presence of mitochondrial impairment and reductive stress **(Fig. 5f-g)**. The ratios were less altered in mutant iPSCs compared to WT iPSCs **(Fig. 5f-g)**. This could be due to the fact that iPSCs are known to predominantly rely on glycolytic metabolism, while the development of NPCs is associated with a shift towards OXPHOS ^37, 38^. In fact, lactate exchange over time remained high only in mutant iPSCs and not in mutant NPCs **(Supplementary Fig. 4f)**. Collectively, these findings suggest that the presence of mHTT causes the elevation of REE, and that these bioenergetic defects could start early during development even before neural commitment.

### Cell-autonomous bioenergetic impairment in pure neurons from HD individuals

We next sought to determine whether functional mitochondrial defects caused by mHTT could occur in human neurons in a cell autonomous manner, independent of potential disruption of other cell types that may be present within cerebral organoids. Several publications investigated iPSC-derived neurons carrying mHTT, but the various differentiation protocols used typically resulted into a heterogeneous population ^20, 34^. Moreover, we wanted to make sure that our findings based on engineered cells reflected phenotypes occurring in patient-derived cells.

To address these aspects, we used three iPSC lines that we recently obtained from individuals affected by HD: HD1 with WT/180Q ^39^ and HD2 and HD3 with WT/58Q and WT/44Q, respectively ^40^ **(Fig. 6a)**. We compared the three HD iPSC lines to three iPSC lines derived from healthy individuals. To obtain a homogenous population of pure neurons, we overexpressed the transcription factor Neurogenin 2 (NGN2) ^41^ in NPCs derived from the three HD iPSC lines and from three healthy control iPSC lines **(Fig. 6b)**. The constructs contained a GFP reporter and allowed derivation of pure neuronal cultures expressing neuronal and synaptic markers **(Supplementary Fig. 5a-b)**. All HD iPSC lines successfully generated NGN2 neurons in a similar manner as control iPSCs **(Supplementary Fig. 5c)**.

**Fig. 6.**
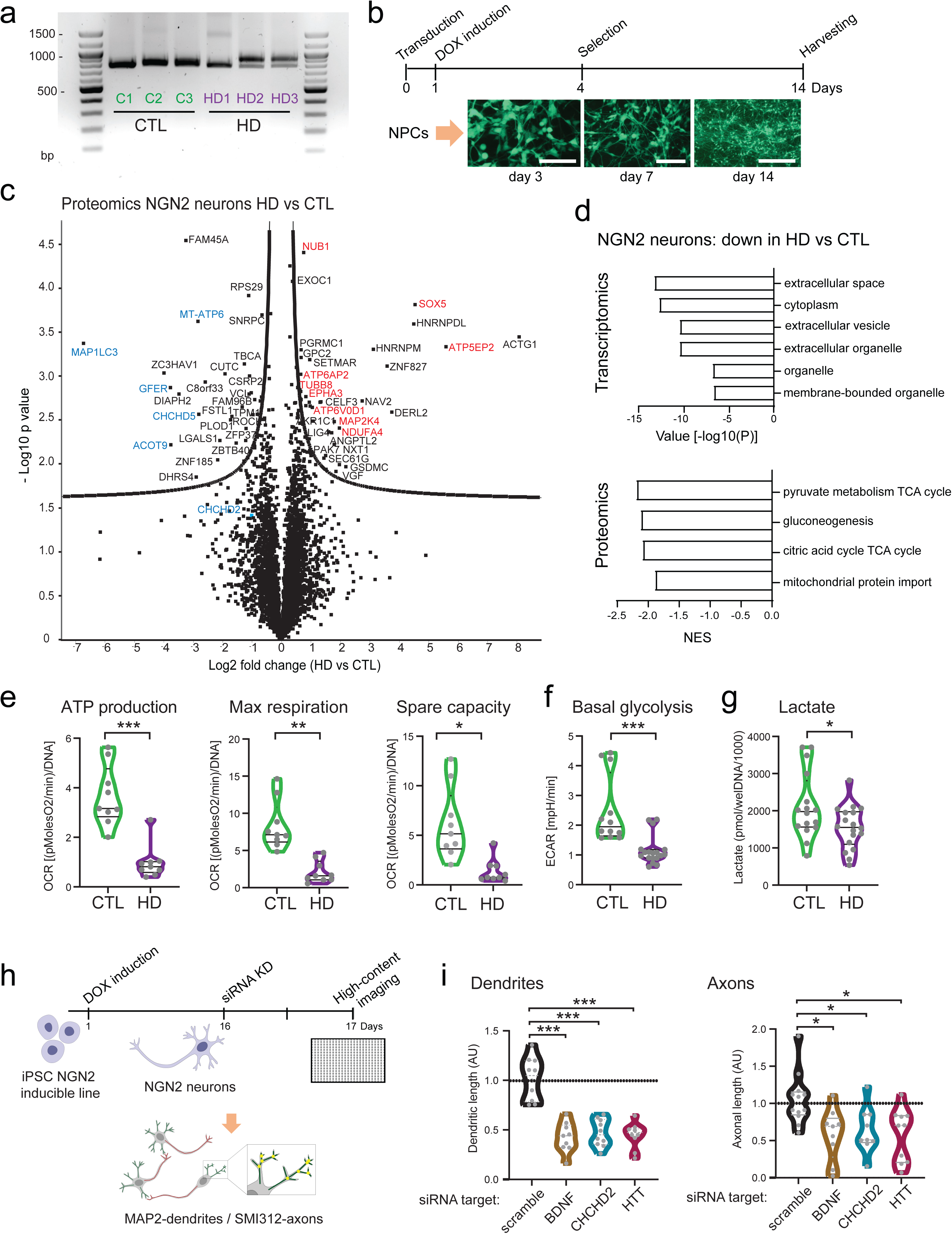
Bioenergetic defects and CHCHD2 knockdown in NGN2 neurons from healthy controls and individuals with HD. **a** PCR analysis of HTT in iPSCs derived from three heathy controls (C1, C2, C3) and three individuals with HD (HD1, HD2, HD3, carrying WT/180Q, WT/58Q, and WT/44Q, respectively). **b** Schematics of Neurogenin 2 (NGN2)-based neuronal induction in NPCs derived from control iPSCs and HD iPSCs. Scale bar: 100 µm. **c** Volcano plot of proteomic datasets in NGN2 neurons derived from control iPSCs (C1, C2, C3) and HD iPSCs (HD1, HD2, HD3) showing statistical significance (log_10_ p-value) versus magnitude of change (log_2_ fold change) (see Supplementary Data 7). Key downregulated proteins in HD neurons are highlighted in blue, key upregulated proteins are highlighted in red. **d** Downregulated pathways identified by either bulk RNA sequencing or proteomics in NGN2 neurons derived from HD iPSCs (HD1, HD2, HD3) compared to NGN2 neurons from control iPSCs (C1, C2, C3) (see Supplementary Data 6-8). **e-f** Mitochondrial bioenergetics based on oxygen consumption rate (OCR) and glycolysis based on extracellular acidification rate (ECAR) measured by Seahorse profiling in NGN2 neurons from control iPSCs (C1, C2, C3) and HD iPSCs (HD1, HD2, HD3). Each dot indicates a biological replicate calculated as the average of all technical replicates. The experiments were performed at least three times. *p<0.05, **p<0.01, ***p<0.001; unpaired two-tailed t test. **g** Quantification of lactate released in the supernatant by NGN2 neurons at the end of Seahorse experiments. **h** Schematics of siRNA knockdown (KD) in NGN2 neurons from NGN2-inducible control iPSCs. Quantification of neuronal arborization was assessed by high-content imaging based on antibodies labeling axons (SMI132) and dendrites (MAP2). **i** Quantification of branching outgrowth in NGN2 neurons following siRNA KD. *p<0.05, ***p<0.001 compared to scramble siRNA KD; Kruskal-Wallis test with Dunn’s multiple comparisons test.

We applied a multi-omics approach to NGN2 neurons derived from the three HD individuals and the three controls. Global transcriptomics and proteomics showed a clear distinction between HD neurons and control neurons **(Supplementary Fig. 5d-e, Supplementary Data 6-9)**. All HD neurons clustered closely even irrespective of different polyQ lengths. These results suggest that the presence of mHTT alters the global expression pattern of human neurons in a cell autonomous manner regardless of the number of CAG repeats **(Supplementary Fig. 5d-e)**.

Downregulated proteins in HD neurons included mitochondrial-related proteins (e.g. GFER, MT-ATP6, ACOT9) **(Fig. 6c)**. Upregulated proteins were also associated with mitochondria (e.g. NDUFA4, ATP5EP2), and furthermore with other processes such as ubiquitin-dependent protein regulation (e.g. NUB, TUBB8) as well as embryonic development and differentiation (e.g. SOX5, MAP2K4, TUBB8) **(Fig. 6c)**. The analysis of the most significant downregulated pathways highlighted mitochondria and energy metabolism in both transcriptomics and proteomics **(Fig. 6d)**. In particular, proteins implicated in mitochondrial import were repressed in the proteome of HD neurons **(Fig. 6d)**. Among those, CHCHD5 protein was significantly downregulated in HD neurons, while the downregulation of CHCHD2 did not reach statistical significance because of its low abundance in differentiated neurons **(Fig. 6c)**.

We next investigated the bioenergetic profiles of NGN2 neurons using the Seahorse technology, which allows probing both mitochondrial and glycolytic metabolism in live cells. In agreement with the omics data, HD neurons exhibited defective mitochondrial function, with diminished ATP production, maximal respiration, and spare respiratory capacity **(Fig. 6e, Supplementary Fig. 5h)**. Unexpectedly, the glycolytic capacity of HD neurons was also compromised, as seen by significantly reduced basal glycolysis **(Fig. 6f)**. In agreement with decreased glycolysis, the amount of lactate in the media of HD neurons was lower than in that of control neurons **(Fig. 6g)**. The global metabolomic measurement resulted in a clear distinction between HD neurons, although not so prominent as transcriptome or proteome **(Supplementary Fig. 5f-g)**. Lactate appeared among the decreased metabolites in HD neurons, together with NAD and acetyl-CoA **(Supplementary Fig. 5g)**. These results suggest that mHTT causes cell autonomous dysregulation of energy metabolism in human neurons, leading to compromised OXPHOS without compensatory activation of glycolysis.

Lastly, we aimed to assess the response of pure neurons to the repression of CHCHD2, given that our data in cerebral organoids indicated that CHCHD2 reduction might be involved in pathogenesis of mHTT. HTT is known to play a role in axonal morphogenesis and function ^7, 42^. Therefore, we focused on neuronal outgrowth capacity of NGN2 neurons using our high-content microscopy based platform **(Fig. 6h)** ^43^. In agreement with the known role of BDNF in neuronal branching ^44^, the downregulation of BDNF in NGN2 neurons impaired the development of axonal and dendritic structures **(Fig. 6i)**. Similar defects in axonal and dendritic branching occurred in NGN2 neurons in which CHCHD2 or HTT were knocked down **(Fig. 6i)**. Since the downregulation of WT HTT also impacted neuronal morphogenesis, therapeutic approaches aiming at repressing HTT may need to be carefully reconsidered. In fact, our model systems carrying mHTT (NPCs, cerebral organoids, neuruloids, and NGN2 neurons) all showed a slight reduction in the amount of WT HTT **(Supplementary Fig. 4g-j)**, implying that HD phenotypes are potentially associated with reduced WT HTT. Taken together, human neurons carrying mHTT develop bioenergetics dysfunctions in a cell autonomous manner, and the repression of CHCHD2 is sufficient to affect their overall capacity of branching outgrowth.

## Discussion

Increasing evidence in mice and humans suggest that mutant HTT could impact the physiological development of the brain ^3, 4, 18^. Our findings based on various human brain organoid differentiation paradigms demonstrates that mHTT disrupts the overall neurodevelopmental process and particularly the organization of neural progenitors. These observations are in agreement with recent data in mice and human fetuses showing that mHTT interferes with the cell cycle of apical progenitors, causing reduced proliferation and premature neuronal lineage specification ^18^. At the same time, the generation of neurons (either appearing physiologically within cerebral organoids or forced through NGN2 overexpression) still occurred in the presence of mHTT.

These results are important for our understanding of the pathomechanisms of HD, as the prevalent view of the disease is that it develops as a degeneration of mature neurons ^45^. If brain defects caused by mHTT instead develop early in life, therapeutic strategies should be administered during development. It is thus possible that therapeutics based on lowering mHTT, as performed in the phase III clinical trial (NCT03761849) developed by Roche ^46^, might not be as effective as hoped for. If such administration is given during adult life, the brain circuity has already been altered and may not be effectively repaired anymore. Accordingly, we found that defects in neural progenitors and brain organoids occurred at a time point where toxic mHTT aggregates have not yet developed. Smaller intracranial volumes can in fact be observed early in pre-manifest HD mutation carriers ^13–15^. These findings raise the possibility that HD pathogenesis may also involve a loss of function mechanism, caused by the reduction of WT HTT. Indeed, WT HTT has been proposed to play a role in neurodevelopment, where it could regulate mitotic spindle orientation thereby determining cortical progenitor fate ^11^, and participate in axonal transport modulation ^47^. In agreement with this, we found that downregulation of WT HTT was sufficient to impair neuronal branching capacity.

By aiming to unveil mechanisms underlying defective human neurodevelopment caused by mHTT, we discovered an early dysregulation of CHCHD2 expression. CHCH domain-containing proteins are imported into the mitochondrion and have been suggested to play a role in neurodegenerative diseases ^30, 31^. CHCHD2 contributes to the maintenance of mitochondrial morphology and dynamics, and its loss has been reported to disrupt mitochondrial organization in models of Parkinson’s disease ^48, 49^. CHCHD2 expression was found to promote neural-ectodermal lineage differentiation of human iPSCs ^50^, indicating that its presence may be important during neurogenesis. Dysregulation of CHCHD2 has also been observed in neural cells carrying mHTT that were differentiated from human embryonic stem cells (hESCs) and iPSCs ^51, 52^. However, the role of CHCHD2 within the pathogenesis of HD remained elusive.

CHCHD2 is known to regulate mitochondrial dynamics and mitochondrial integrated stress response (mISR) ^25^. mISR is a process that rises in stress conditions such as those associated with impaired OXPHOS ^33^. Accordingly, we identified a mISR signature in human neural cells carrying mHTT that was associated with aberrant mitochondrial morpho-dynamics. Importantly, elimination of the poly-Q/poly-P region reverted the abnormal CHCHD2 expression and the associated mISR signature and mitochondrial defects. Hence, mHTT leads to CHCHD2 dysregulation, which in turn impairs mitochondrial function and organization.

Dysfunction of mitochondrial activity and network morphology have been observed in multiple model systems of HD ^34, 53–55^. Even Pridopidine, currently being investigated as a treatment for HD in Phase III study (NCT04556656), has been shown to repress mitochondrial stress and network defects ^56^. However, again in agreement with a neurodegenerative view of the disease, mitochondrial dysfunction in HD models is usually thought to occur mainly within mature neuronal cells. Our data instead indicate that aberrant CHCHD2 expression and related mitochondrial morpho-function impairment can occur early during neurodevelopment. Such early mitochondrial dysfunctions is expected to affect the overall process of neurodevelopment ^57^. Impairment of the cellular NAD+/NADH redox balance caused by metabolic perturbation can in fact lower the global rate of protein synthesis, thereby slowing down the segmentation clock and overall developmental rate ^58^. In particular, mitochondrial morphology and metabolism are emerging as important regulators of early neurogenesis ^59–61^. Mitochondrial dynamics within NPCs regulate the pace of development ^62^, and help orchestrating the balance between proliferation and neuronal specification ^63^. A correct metabolic programming towards OXPHOS starts to develop at the level of NPCs ^37, 38, 64, 65^. In our models, we found that both undifferentiated iPSCs and NPCs were able to respond to OXPHOS impairment by upregulating glycolytic metabolism. However, an impaired NAD/NADH ratio became evident in NPCs. Differentiated neurons could not compensate these defects and developed cell autonomous impairment of both OXPHOS and glycolysis. The establishment of neural fate might then pose greater pressure on cells that start relying more on OXPHOS, ultimately leading to metabolic stress and functional impairment. Our data thus indicate that mHTT interferes with the physiological metabolic programming, which is required for enabling physiological neural fate commitment.

Individuals with HD often suffer from weight loss and cachexia irrespective of food intake ^66–68^. In fact, the metabolic network impairment has been suggested as a progression biomarker of pre-manifest HD ^69^. We found that mHTT caused a metabolic state indicative of elevated energy consumption at rest. This feature, also known as hypermetabolism, leads to weight loss, and is associated with greater functional decline in cancer patients or individuals with amyotrophic lateral sclerosis ^70^. Hypermetabolism has recently been found to occur in the presence of OXPHOS-impairing mutations that can lead to mISR activation ^33^. Previously, we and others have demonstrated that OXPHOS-impairing mutations cause aberrant neuronal morphogenesis and defective brain organoid development ^71, 72^. In agreement with these data, we found that CHCHD2 reduction in pure neurons impaired neuronal branching capacity. Hence, it is possible that CHCHD2 could represent a target for early intervention in HD, as its modulation would impact mitochondrial morpho-function, which in turn could be important for morphogenesis and neurodevelopment on the one hand and for overall systemic metabolic resilience on the other.

It is important to highlight that the cells carrying 70Q/70Q represent an extreme pathological condition in whose allelic combination the compensation effect of the WT allele is not present. In fact, the application of 70Q/70Q genotype was intended to “fast-forward” the disease phenotype as it allows the clear transcriptional dissection of mHTT. Accordingly, WT/70Q cells not always fully mimicked the defects observed for 70Q/70Q cells. At the same time, it was reassuring to see that pure neurons from HD patients carrying WT/180Q, WT/58Q, and WT/44Q recapitulated similar defects regardless of the poly-Q length. Indeed, a recent study using cerebral organoids obtained from hESCs carrying WT/45Q, WT/65Q and WT/81Q observed described similar features indicative of neurodevelopment impairment ^23^. Nonetheless, additional studies focusing on various HD forms are warranted for further elucidating the importance of CHCHD2 dysregulation in the pathogenesis of HD. Further investigations are also needed to address whether somatic deletion of the polyQ region in post-mitotic neurons, in contrast to applied in in-frame deletion in iPSCs, could be beneficial for individuals with HD to possibly reverse disease features. Moreover, it remains to be determined whether the transcription factor CHCHD2 could represent a direct binding partner of mHTT.

## Author contributions

Conceptualization, A.P., P.L.; Methodology, P.L., S.L., B.M., A.R-W., S.L., C.M., W.D., Y.R., L.A.M.K., H.W., D.M., M.O., N.N., T.H., A.B.; Formal Analysis, D.O., P.R., P.G., I.L.; Resources, A.P. E.E.W; N.R.; R.K., J.K., J.P., E.M., S.C., H.O., J.M.; Writing –Original Draft, A.P.; Writing – Review & Editing, A.P., J.J.M., J.P., S.C., A.R-W.; Supervision, A.P., J.J.M., J.K., S.C., N.R., H.O., E.E.W., J.P.; Visualization, A.P., S.L.; Funding Acquisition, A.P., S.C., J.J.M., H.O., J. P.

## Acknowledgements

We are grateful to Daria Mochly-Rosen for help with the mitochondrial network morphology assay. We acknowledge support from the Deutsche Forschungsgemeinschaft (DFG) (PR1527/5-1 and PR1527/6-1 to A.P., RTG 2155 ProMoAge to H.O. and L.A.M.K., SFB167 B07 to J.P.), the Berlin Institute of Health (BIH) (to S.D., J.P., R.K., and A.P.), the Bundesministerium für Bildung und Forschung (BMBF) (AZ. 031L0211 and 01GM2002A to A.P.), the Medical Faculty of Heinrich Heine University (FoKo grant to A.P. and S.C.), and The National Science Centre, Poland (NCN grant No. 20 16/22/M/NZ2/00548 and 2017/27/B/NZ1/02401 to P.L.). We acknowledge the Center for Advanced Imaging (CAi) at Heinrich Heine University Düsseldorf for providing access to the PerkinElmer Operetta CLS (DFG grant number INST 208/760-1 FUGG) and Olympus FV3000 microscope.

## Msaterial and methods

### Human iPSC lines and culture

WT/WT iPSCs for genome editing were derived using Sendai viruses from a healthy middle-aged male individual (BIHi050-A/SCVI113). iPSCs from three male individuals with HD were described before: HD1 with WT/180Q (BIHi035-A) ^39^, HD2 with WT/58Q (BIHi288-A) ^40^, and HD3 with WT/44Q (BIHi033-A) ^40^. Control iPSCs used for comparison with HD iPSCs were derived before: C1 (TFBJ, HHUUKDi009-A) ^37^, C2 (XM001, BIHi043-A) ^73^, and C3 (BIHi005-A). For engineering an inducible NGN2 iPSC line, we used the control healthy iPSC line C3 (BIHi005-A). All iPSCs were cultured feeder-free on Matrigel-coated (Corning, USA) 6-well plates (Greiner Bio-One, Austria) in iPS-Brew consisting of StemMACS iPS-Brew XF basal medium supplemented with 1:50 StemMACS iPS-Brew XF supplement (both Miltenyi Biotec, Germany) and 1:500 MycoZap-Plus-CL (Lonza, Switzerland). iPSC cultures were kept in a humidified atmosphere of 5% CO_2_ and 5% oxygen at 37°C. We routinely monitored against mycoplasma contamination using PCR. Cells were passaged at 70-80% confluence with 0.5 µM EDTA (Invitrogen, USA) in 1xPBS (Gibco, USA). 10 µM Rock inhibitor (Enzo Biochem, USA) was added after splitting to promote survival. Ethical approval for using iPSCs from HD individuals and control subjects was obtained from the Ethic Committee of the University Clinic Düsseldorf (study number 2019-681 approved on October 11, 2019).

### Engineering Huntingtin in human iPSCs

Isogenic clones with a homozygous (HTTWT/70Q) and heterozygous (HTT70Q/70Q) allelic combinations carrying 70Q insert were engineered from control (WT/WT) BIH-SCVI113 iPSC line with heathy background. We used CRISPR/Cas9 induced homology directed repair (HDR) with 2354 nts dsDNA donor template encoding patient reference sequence carrying 70Q consisting of both CAG and CAA triplets including 3’ and 5’ homology arms. Constructs were commercially synthetized as a synthetic HTT donor construct and assembled from synthetic oligonucleotides by GeneArt Gene Synthesis service (Thermo Fisher Scientific, USA). Silent mutations within single guide RNAs (sgRNAs) were introduced in close proximity to the each of two sgRNA protospacer adjacent motif (PAM) sites (3 and 6 nt downstream) to prevent recurrent Cas9 cutting in edited cells. Synthesized template was inserted into pMK-RQ (kanR)/pMA-RQ (ampR) plasmid, purified from transformed bacteria, determined by spectroscopy and verified by sequencing. sgRNAs (sgHTT#1, sgHTT#2) targeting upstream and downstream of the HTT polyQ region were designed using CRISPOR (http://crispor.tefor.net/) **(Supplementary Fig. 1a, Supplementary Data 10)**. The oligomer pairs were annealed and cloned separately into two pU6-CAG-eCas9-Venus plasmids carrying eSpCas9 variant from eSpCas9(1.1) plasmid (Addgene ID 71814) to reduce off-target effects and improve on-target cleavage ^26^. To upregulate homology directed repair of eCas9 induced DSBs with dsDNA donor template, we applied ectopic expression of BRCA1 and BCL-XL. To downregulate the non-homologous end joining (NHEJ) pathway we applied dominant-negative subfragment of 53BP1 (dn53BP1), which counteracts endogenous 53BP1 ^27^. Plasmids encoding components of the DNA repair pathways (human BRCA1, BCL-XL and mouse dn53BP1) were kindly obtained from Bruna Paulsen (dn53BP1) and Xiao-Bing Zhang (BCL-XL). For generation of dn53BP1, a fragment containing the tudor domain (residues 1,221 to 1,718 of mouse 53BP1) was amplified and sub-cloned into a CAG expression plasmid and sequenced. For the generation of BCL-XL, a fragment containing the BCL-XL was amplified sub-cloned into a pEF1-BFP expression plasmid and sequenced. pU6-sgHTT#1-CAG-eCas9- Venus and pU6-sgHTT#2-CAG-eCas9-Venus plasmids targeting upstream and downstream of the CAG/CCG stretches of the HTT gene, plasmids encoding 70Q HDR template and BRCA1, BCL-XL and mouse dn53BP1 were transformed into super competent DHA bacteria strain using heat shock and cloned. Plasmid DNA was extracted using the PureYield plasmid miniprep system (Promega) and transfected into control the iPSC line that reached at least 80% confluency of the 6-well cell culture plate format. In addition to the BIH-SCVI113 hiPSCs with introduced 70Q templates, we also generated the clones harboring an HTT alleles underwent NHEJ mediated excision of the poly Q/P repeats region by in frame reannealing of the DSB. By transfection of pU6-sgHTT#1-CAG-eCas9-Venus and pU6-sgHTT#2-CAG- eCas9-Venus with BCL-XL plasmids, we generated DSBs which upon re-ligation with the lack of HDR template resulted into an in-frame HTT coding region, lacking the N-terminal Q/P repeats. Transient transfection of plasmids was carried out in BIH-SCVI113 iPSC line grown in feeder-free conditions in StemMACS™ iPS-Brew XF culture media (Miltenyi Biotec, Germany) in a 6-cell culture plate. One day prior to transfection, we dissociated the cells using Accutase (Sigma-Aldrich, USA) and seeded ∼ 1 × 10^5^ cells per well of a pre-coated 6-well plate as single cells or small clumps. Cell were cultivated in fresh medium containing 10 mM Rock inhibitor (Enzo Biochem, USA) overnight. Transfection was performed using Lipofectamine™ 3000 Transfection Reagent (Thermo Fisher Scientific, USA), according to the manufacturer’s protocol. The plasmids were diluted up to 2 mg DNA in 125 ml of Opti-MEM reduced serum medium and added as the DNA-lipid complex to one well of a 6-well plate in a dropwise manner with addition of 5 mM Rock inhibitor to the culture medium for 24 h. Medium was changed on the following day and the cells were kept 48 h in culture until fluorescence-activated cell sorting (FACS). Dissociated cells using Accutase for 5 min were washed and resuspended with PBS. Then, cells were filtered using Falcon polystyrene test tubes (Corning, USA) and transferred to Falcon polypropylene test tubes (Corning, USA). Two colors FACS enrichment was performed using BD FACSAria III at the MDC FACS Facility for cells expressing high levels of eCas9-sgRNAs, and BCL-XL (Venus, BFP). Sorted cells were suspended in recovery mTeSR™ medium (StemCell Technologies, Canada) with 1X Penicillin-Streptomycin (P/S) (Gemini Bio-products) and ROCK inhibitor and plated in low concentrations onto 6-well plates for the establishment of single cell derived colonies (5,000 cells/well). Growing single cell-derived colonies were transferred from 6-well plates to one well each of 24-well plate and maintained until the colony grew big enough to be partially harvested for DNA isolation using Phire Animal Tissue Direct PCR Kit (Thermo Fisher Scientific, USA) according to manufacturer’s protocol. PCR reaction was carried out using 10 ng gDNA in 25 ml with chemical hot-Start AmpliTaq Gold DNA Polymerase (Thermo Fisher Scientific, USA) with annealing temperature 55°C and GC enhancer reagent (Thermo Fisher Scientific, USA). For Sanger sequencing the PCR products are gel purified using e.g. the Wizard SV Gel and PCR Clean-Up System (Promega) and cloned into pJet cloning vector using CloneJET PCR Cloning Kit (Thermo Fisher Scientific, USA). Cloned PCR products were submitted to LGC (https://www.lgcgroup.com) for Sanger sequencing. Karyotype analysis was performed by MDC Stem Cell Core Facility. Briefly, DNA was isolated using the DNeasy blood and tissue kit (Qiagen, USA). SNP karyotyping was assessed using the Infinium OmniExpressExome-8 Kit and the iScan system from Illumina. CNV and SNP visualization were performed using KaryoStudio v1.4 (Illumina). Primer sequences, gRNA sequences, and HDR sequence are reported in **Supplementary Data 10**.

### Derivation of neural progenitor cells (NPCs)

To generate neural progenitor cells (NPCs) from iPSCs, we applied our published protocol ^74^. Briefly, iPSCs were detached from Matrigel-coated plates using Accutase (Sigma-Aldrich, USA) and the collected sedimented cells were transferred into low-attachment petri dishes and kept for two days in M1 medium (1x KnockOut DMEM, 1x KnockOut serum, 0.1 mg/mL PenStrep, 2 mM Glutamine, 1x NEAA, 1 mM Pyruvate [all Gibco, USA], 1x MycoZap-Plus-CL [Lonza, Switzerland], 3 µM CHIR 99021 [Caymen Chemical, USA], 10 µM SB-431542 [Caymen Chemical, USA], 1 µM Dorsomorphin [Sigma-Aldrich, USA], 500 nM Purmorphamine [Miltenyi Biotec, Germany]). From day 2 to day 6, the media was switched to M2 medium (50% DMEM-F12, 50% Neurobasal, 0.5x N-2 supplement, 0.5x B-27 supplement without Vitamin A, 0.1 mg/mL PenStrep, 2 mM Glutamine [all Gibco, USA], 1x MycoZap-Plus-CL [Lonza, Switzerland], 3 µM CHIR 99021 [Caymen Chemical, USA], 10 µM SB-431542 [Caymen Chemical, USA], 1 µM Dorsomorphin [Sigma-Aldrich, USA], 500 nM Purmorphamine [Miltenyi Biotec, Germany]). On day 6, the suspended cells were transferred onto Matrigel-coated (Corning, USA) 6-well plates (Greiner Bio-One, Austria) using sm+ medium (50% DMEM-F12, 50% Neurobasal, 0.5x N-2 supplement, 0.5x B-27 supplement without Vitamin A, 0.1 mg/mL PenStrep, 2 mM Glutamine [all Gibco, USA], 1x MycoZap-Plus-CL [Lonza, Switzerland], 3 µM CHIR 99021 [Caymen Chemical, USA], 500 nM Purmorphamine [Miltenyi Biotec, Germany], 150 µM ascorbic acid [Sigma-Aldrich, USA]). NPCs were maintained in this media without rock inhibitor and used for experiments between passage 7 and 20.

### Generation of unguided cerebral organoids

Unguided cerebral organoids were generated from WT/WT and 70Q/70Q iPSCs using a modified version of a published protocol ^75^. At day 0, iPSCs from one 80% confluent well were washed with 1x PBS (Gibco, USA) and treated with Accutase (Sigma-Aldrich, USA) for 3 min at 37°C. Subsequently, cells were collected in 5 ml iPS-Brew and spun down for 5 min at 270 x g. After removing the supernatant, cells were resuspended in 1 ml embryoid body (EB) media (1x DMEM-F12, 1:5 KnockOut-Serum, 1x NEAA, 0.1 mg/mL PenStrep, 1x GlutaMAX [all Gibco, USA], 7 nM 2-mercaptoethanol [Merck, Germany], 4 ng/mL bFGF [PeptroTech, USA], 10 µM Rock inhibitor [Enzo Biochem, USA]). Cells were counted manually and 9,000 cells were plated in 150 µL EB media in each well of a round bottom, ultra-low attachment 96-well plate (Corning, USA). Next, the plate was spun down at 500 rpm for 2 min to assure the aggregation of cells in the center of the wells. At day 2 and day 4, 50% of the EB media was replaced with EB media without ROCK inhibitor and bFGF. At day 5, each EB was transferred to a well of a 24-well plate (Greiner bio-one, USA) in 250 µL neural induction media (IM) (1x Neurobasal, 1x N-2 supplement, 1x NEAA, 1x GlutaMax, 0.1 mg/mL PenStrep [all Gibco, USA], 1 µg/mL heparin [Sigma-Aldrich, USA]). At day 6 and 10, another 250 µL IM was added to each well. At day 11, organoids were embedded in droplets of 30 µL Matrigel (Corning, USA) as previously described ^75^, and 16 organoids each were transferred to 60×15 mm cell culture dishes (Greiner bio-one, USA) and cultured for 48 h in 5 mL IM at 37°C with 5% CO_2_. After this, IM was replaced with neuronal differentiation media 1 (DM1) (50% DMEM-F12, 50% Neurobasal, 0.5x N-2 supplement, 2x B-27 supplement without vitamin A, 0.1 mg/mL PenStrep, 1x GlutaMax, 0.5x NEAA [all Gibco, USA], 0.35 nM 2-mercaptoethanol [Merck, Germany], 2.5 µM insulin [Sigma-Aldrich, USA], 3µM CHIR99021 [Caymen Chemical, Germany]). After 2 days, the media was replaced by fresh DM1 and at day 17 the organoids were placed on an orbital shaker (85 rpm) and the media was replaced by DM2 (50% DMEM- F12, 50% Neurobasal, 0.5x N-2 supplement, 1x B-27 supplement, 0.1 mg/mL PenStrep, 1x GlutaMax, 0.5x NEAA, 1:100 chemically defined lipid concentrate [all Gibco, USA], 0.35 nM, 2-mercaptoethanol [Merck, Germany], 2.5 µM insulin [Sigma-Aldrich, USA]) . After 3 days, with a full media refreshment after 2 days, the media was replaced with DM3 (50% DMEM/F12, 50% Neurobasal, 0.5x N-2 supplement, 1x B-27 supplement, 0.1 mg/mL PenStrep, 1x GlutaMax, 0.5x NEAA, 1:100 chemically defined lipid concentrate [all Gibco, USA], 0.35 nM 2- mercaptoethanol [Merck, Germany], 2.5 µM insulin [Sigma-Aldrich, USA], 0.4 µM ascorbic acid [Sigma-Aldrich, USA], 12 mM HEPES solution [Biochrom, Germany]. From this point on, full media replacements took place every 3-4 days. At day 40, the media was changed to maturation media (MM) (1x Neurobasal, 1x B-27 supplement, 0.1 mg/mL PenStrep [all Gibco, USA], 0.35 nM 2-mercaptoethanol [Merck, Germany], 0.4 µM ascorbic acid [Sigma-Aldrich, USA], 20 ng/mL BDNF [Miltenyi Biotec, Germany], 20 ng/mL GDNF [Miltenyi Biotec, Germany], 0.5 mM cAMP [Sigma-Aldrich, USA], 12 mM HEPES solution [Biochrom, Germany]). Media was checked for mycoplasma routinely. Organoid size was measured using ImageJ software.

### Generation of guided region-specific brain organoids

Cortical organoids were generated following a protocol previously published ^76^ with minor modifications. On day 0, iPSCs at 80% confluence were incubated with Accutase (Sigma-Aldrich, USA) at 37°C for 7 min and dissociated into single cells. For aggregation into spheroids, approximately 3 × 10^6^ single cells were seeded per AggreWell-800 well (StemCell Technologies, Canada) in StemMACS iPS Brew XF (Miltenyi Biotec, Germany) supplemented with 10 μM ROCK inhibitor (Enzo Biochem, USA), centrifuged at 100 x g for 3 min, and then incubated at 37°C in 5 % CO_2_. After 24 hours, spheroids consisting of approximately 10,000 cells were collected from each microwell by gently pipetting the medium up and down with a cut P1000 pipet tip and transferred into a petri dish coated with Anti-adherence solution (StemCell Technologies, Canada). From day 1-6, ES Medium (KnockOut DMEM, 20% Knockout Serum Replacement, 1x NEAA, 1x Sodium Pyruvate, 1x GlutaMAX [all Gibco, USA], 1x MycoZap-Plus-CL [Lonza, Switzerland], 2.5 μM dorsomorphin [Sigma-Aldrich, USA], 10 μM SB-431542 [Caymen Chemical, USA]) was changed daily. From day 7-22, neural medium (Neurobasal-A, B-27 Supplement without Vitamin A, 1x Glutamax [all Gibco, USA], 1x MycoZap-Plus-CL [Lonza, Switzerland]) supplemented with 20 ng/mL EGF (R&D Systems, USA), and 20 ng/mL FGF2 (R&D Systems, USA). From day 22, the neural medium was supplemented with 20 ng/mL BDNF (Miltenyi Biotec, Germany), 20 ng/mL NT-3 (PeproTech, USA), 200 µM ascorbic acid (Sigma-Aldrich, USA), 50 µM db c-AMP (StemCell Technologies, Canada) and 10 µM DHA (MilliporeSigma, USA). From day 46, only neural medium containing B-27 Plus Supplement (Gibco, USA) was used for media changes every 2-3 days.

We generated NPC-derived midbrain organoids according to a previously published protocol, with some modifications ^29^. On day 0, we detached NPCs using Accutase (Sigma-Aldrich, USA) to obtain a single cell suspension. Using an automatic cell counter (Cytosmart, Netherlands) we gated the cell size 5-19 µM and seeded 9,000 cells per well onto a low-attachment U-bottom 96-well plate (FaCellitate, Germany) in 150µL of basal media (50% DMEM-F12, 50% Neurobasal, 0.5x N2 supplement, 0.5x B-27 supplement without Vitamin A, 1x GlutaMAX [all Gibco, USA], 1x MycoZap-Plus-CL [Lonza, Switzerland]) supplemented with 0.5 µM PMA (Miltenyi Biotec, Germany), 3 µM CHIR 99021 (Sigma- Aldrich, USA), and 100 µM ascorbic acid (Sigma-Aldrich, USA). After 2 days, we started ventral patterning for 4 days (in two feedings) by supplementing the basal media with 100 µM ascorbic acid, 1 µM PMA, 1ng/mL BDNF (Miltenyi Biotec, Germany) and 1ng/mL GDNF (R&D Systems, USA). On day 6 we switched to maturation media by supplementing the basal media with 100 µM ascorbic acid, 2 ng/mL BDNF, 2 ng/mL GDNF, 1 ng/mL TGF-β3 (StemCell Technologies, Canada), 100 µM db c-AMP (StemCell Technologies, Canada) and the addition of 5 ng/mL Activin A (StemCell Technologies, Canada). From day 6 on, we refreshed the maturation media (without Activin A, which is exclusively added on day 6) 3 times a week. To measure organoid size, we first took pictures of the organoids every other day with a Nikon Eclipse Ts2 inverted routine microscope using a 4x objective. Subsequently we analyzed the images with ImageJ software by manually drawing a circle around the organoid, as previously described ^71^. We measured the area and perimeter from at least 5 organoids per cell line for each time point.

### Engineering of an inducible NGN2 iPSC line

We used the control healthy iPSC line C3 (BIHi005-A) to engineer a doxycycline inducible NGN2 expression cassette to obtain the line BIHi005-A-24. To generate this cell line, we used the strategy of TALEN based gene-editing. Each AAVS1 locus was changed differently; one AAVS1 allele inserted the constitutive expression cassette of the reverse trans activator domain (m2rtTA) and the other one the mNgn2-P2A-GFP-T2A-Puromycine cDNA sequence under the tetracycline responsive element (TRE). To generate the AAVS1-TRE-mNgn2-P2A- GFP-T2A-Puromycine donor plasmid, we amplified the TRE-mNgn2-P2A-GFP-T2A- Puromycine sequence from LV-NEP (YS-TetO-FUW-Ng2-P2A-EGFP-T2A-Puro) plasmid from Dr. Thomas Sudhof lab ^41^, and cloned it into the plasmid AAVS1-iCAG-copGFP (Addgene 66577). To make the lines inducible for doxycycline, the reverse TET trans activator (m2rtTA) was inserted on the other allele of the AAVS1 locus using the m2rtTA plasmid containing a Neomycin resistance gene (addgene 60843). The TALEN plasmids to target AAVS1 locus were used from addgene: hAAVS1-TAL-L (35431) and hAAVS1-TAL-R (35432). BIHi005-A iPSC cells were transfected with TALEN plasmids targeting AAVS1 locus and donor plasmids (AAVS1-TRE-mNgn2-P2A-GFP-T2A-Puro and AAVS1-NEO-M2rtTA). Then we selected the clones which were resistant for both puromycin and neomycin antibiotics. Next, we derived single cell clones as described ^77^, and selected the clones that express GFP and differentiated to neurons upon exposure to 3 µg/mL doxycycline. The selected clones showed typical pluripotent stem cell morphology, expressed pluripotent markers and showed normal karyotype.

### Derivation of NGN2 neurons

NGN2 neurons were generated from NPCs from control iPSC lines C1, C2, C3, and from HD patient-derived iPSC lines H1, H2, H3 by modifying a published protocol ^41^. For NGN2 virus production, HEK 293 cells were seeded at 70% confluency in DMEM medium in a 150 cm^2^ dish. After cell attachment, the medium was replaced and supplemented with 25 µM Chloroquine and the cells were transfected according to the manufacturer’s instructions using Lipofectamine 2000 (Thermo Fisher Scientific, USA) and the following plasmid mix: 8.1 µg pMD2.G, 12.2 µg pMDLg-pRRE, 5.4 µg pRSV-Rev, 5.4 µg FUW-M2-rtTA, 14.3 µg TetO-FUW-NGN2, 14.3 µg TetO-FUW-EGFP. After 24 h, the medium was replaced with DMEM-F12 (Gibco, USA) and 10 µM sodium butyrate (Sigma-Aldrich, USA). Virus-containing supernatant was collected 24 h and 48 h later. To concentrate the virus particles, the supernatant was first centrifuged at 500 x g for 10 min to remove cells and debris and then mixed with 1 volume of cold Lenti-X concentrator (Takara Bio, Japan) to every 3 volumes of lentivirus-containing supernatant and kept overnight at 4°C. Afterwards, the mixture was centrifuged at 1,500 x g for 45 min at 4°C, and the resulting pellet was resuspended in PBS (Gibco, USA), 1:100 of the original volume. Virus aliquots were stored at -80°C until further use. For NGN2 neuron generation and culture, 2.5 × 10^6^ NPCs per well were seeded on a Matrigel-coated (Corning, USA) 6-well plate (Greiner Bio-One, USA) using sm+ medium (see NPC generation for media composition). After cells attached (2 h – 24 h), the medium was replaced and supplemented with 4 µg/mL polybrene and 2.25 × 10^6^ transducing units of both EGFP and NGN2 lentivirus. On the next day (day 0), the cells were washed 3 times with PBS and 3 volumes sm+medium mixed with 1 volume NGN2 medium (Neurobasal, 1x B-27 supplement, 1x NEAA, 2 mM Glutamine, 0.1 mg/mL Pen/Strep [all Gibco, USA], 1x MycoZap-Plus-CL [Lonza, Switzerland], 10 ng/mL human NT-3 [PeproTech,USA], 10 ng/mL human BDNF [R&D Systems, USA]) supplemented with 2 µg/mL doxycycline (Sigma-Aldrich, USA). On day 3, medium was changed to 100% NGN2 medium supplemented with 0.8 µg/mL puromycin (Sigma-Aldrich, USA). On day 5, selection process was stopped by replacing medium without antibiotics. Medium was changed every other day until day 14, where the cells were harvested for subsequent experiments.

For NGN2 neuron generation based on the inducible NGN2 iPSC line BIHi005-A-24, 3 × 10^5^ iPSCs per well were seeded in a Geltrex (Gibco, USA) coated 6-well plate (Greiner Bio-One, Austria) using StemMACS iPS-Brew medium supplemented with 10 µM Rock inhibitor (Enzo Biochem, USA). On day 0 and day 1, we added induction medium (DMEM-F12, 1x N2 supplement, 1x NEAA, 1 mg/mL Pen/Strep [all Gibco, USA], 10 ng/mL human NT-3 [PeproTech, USA], 10 ng/mL human BDNF [Miltenyi Biotec, Germany], 0.2 µg/mL murine laminin [Sigma-Aldrich, USA]) freshly supplemented with 3 µg/mL doxycycline (Sigma-Aldrich, USA). On day 2, medium was changed to neuronal medium (Neurobasal, 1x B-27 supplement, 1x GlutaMAX, 1 mg/mL Pen/Strep [all Gibco, USA], 1x MycoZapTM Plus-CL (Lonza, Switzerland), 10 ng/mL human NT-3 [PeproTech, USA], 10 ng/mL human BDNF [Miltenyi Biotec, Germany], 0.2 µg/mL murine laminin [Sigma-Aldrich, USA]) freshly supplemented with 3 µg/mL doxycycline (Sigma-Aldrich, USA). Medium was exchanged daily until day 4. From day 6 on, half of the medium was replaced every other day with neuronal medium freshly supplemented with 3 µg/mL doxycycline (Sigma-Aldrich, USA) and 2 µM AraC (Sigma-Aldrich, USA) until the cells were harvested for subsequent experiments.

### siRNA-based knock-down experiments and branching analysis

For siRNA-based knock-down experiments, siRNA (Dharmacon, USA) was incubated with JetPRIME transfection reagent (Polyplus-transfection, France) and µClear black 384-well-plates (Greiner Bio-One, Austria) were coated with 1 pmol siRNA per well. The plates were dried in a SpeedVac and stored at 4°C wrapped with parafilm until use. Before adding the neurons on the plates, they were additionally coated for 1 h at 37°C with ready-to-use Geltrex (Gibco, USA). 13 days-old neurons were detached with Accutase (Sigma-Aldrich, USA) for 3 min and collected. The cells were centrifuged at 100 x g for 5 min and resuspended in neuronal medium. We quantified neuronal branching as we previously described ^43^. Briefly, 5,000 neurons were seeded per well to the 384-well plate and cultured for 4 days. At day 17 of differentiation, the neurons were fixed by adding 8% PFA (Thermo Fisher Scientific, USA) to the medium and incubated for 20 min. After washing 3 times with PBS (Gibco, USA), 0.05% sodium azide (Sigma-Aldrich, USA) in PBS solution was added to the wells and the plate was stored at 4°C wrapped with parafilm until staining. For staining, the cells were blocked for 1 h at RT with blocking solution (PBS [Gibco, USA], 3% BSA, 0.05% sodium azide, 0.5% Triton-X-100 [all Sigma-Aldrich, USA]). Then, they were treated with primary antibodies guinea pig anti-MAP2 (1:1,000; Synaptic Systems, Germany) and mouse anti-SMI312 (1:500; BioLegend, USA) in blocking solution over night at 4°C wrapped with parafilm. Afterwards, they were washed 3 times with PBS and treated with secondary antibodies against guinea pig (AF 488, 1:500; Sigma-Aldrich, USA) and mouse (AF 568, 1:500; Thermo Fisher Scientific, USA) and Hoechst 33342 (1:2500; Life Technologies, USA) in blocking solution for 1 h at RT. After 3 washes with PBS, the wells were filled with 0.05% sodium azide in PBS and stored at 4°C wrapped with parafilm. They were imaged with the high-content microscope Operetta (PerkinElmer, USA) with a 0.4 NA 20X air objective and 25 images per well. The images were analyzed with the open-source software CellProfiler regarding their axon and dendritic growth and arborization. The pipeline can be found here: https://github.com/StemCellMetab/neurite-outgrowth. The data was summarized by calculating the median per well and to compare different plates, the numbers were compared to the median of the control wells (non-targeting scrambled siRNA). siRNA sequences are reported in **Supplementary Data 10.**

### Total RNA sequencing

Bulk RNA-sequencing was carried out for the following samples (each done in biological triplicates) Dataset 1: WT/WT iPSCs, NPCs, d28 cerebral organoids, and d49 cerebral organoids, 70Q/70Q iPSCs, NPCs, d28 cerebral organoids, and d49 cerebral organoids, WT/70Q NPCs. Dataset 2: NGN2 neurons from C1, C2, C3, HD1, HD2, HD3. Total RNA was isolated using the Qiagen isolation kit and quality-checked by Nanodrop analysis (Nanodrop Technologies). Briefly, total RNA was mixed with 1 μg of a DNA oligonucleotide pool comprising 50-nt long oligonucleotide mix covering the reverse complement of the entire length of each rRNA (28S rRNA, 18S rRNA, 16S rRNA, 5.8S rRNA, 5S rRNA, 12S rRNA), incubated with 1U of RNase H (Hybridase Thermostable RNase H, Epicentre), purified using RNA Cleanup XP beads (Agencourt), DNase treated using TURBO DNase rigorous treatment protocol (Thermo Fisher Scientific) and purified again with RNA Cleanup XP beads. rRNA-depleted RNA samples were further fragmented and processed into strand-specific cDNA libraries using TruSeq Stranded Total LT Sample Prep Kit (Illumina) and sequenced on NextSeq 500, High Output Kit, 1x 150 cycles. Raw sequencing reads were mapped to the human genome (GRCh38 assembly) using STAR (version 2.6.0c) aligner ^78^. We used the default settings, with the exception of --outFilterMismatchNoverLmax, which was set to 0.05. Reads were counted using the htseq-count tool, version 0.9.1 ^79^, with gene annotation from GENCODE release 27 ^80^. Differential gene expression analysis was performed using the DESeq2 (version 1.20.00) R package ^81^. All genes with the adjusted P-value lower than 0.05 were considered differentially expressed. Functional enrichment analysis was done using the gProfileR R package ^82^, version 0.6.6, with default settings. All expressed genes were used as background. For heatmap generation for pathway genes of Dataset 1, bulk-RNAseq count matrices were loaded into R (v. 4.2.2). Additional organoid data from neuruloids ^24^ was processed from sc-RNAseq data to pseudobulk data by randomly pooling single cells into 3 bulk replicates per cell line per condition (HD and WT/WT) and then also loaded into R. The raw matrices were then normalized to counts per millions (CPM) and filtered to only contain genes with more than 1 CPM using edgeR (v 3.40.2). For the genes corresponding to a pathway of interest, all the different samples and their corresponding control samples were compared with edgeR, yielding the calculated log2fc and adjusted p values. For each pathway the obtained results were plotted in a separate heatmap with colors indicating the magnitude of the log2fc and stars indicating the statistical significance. The case of 0.05 > p_val > 0.01 corresponds to one star, 0.01 > p_val > 0.001 corresponds to two stars, and everything below corresponds to three stars. All R scripts are available on request. RNA-seq datasets are deposited into NCBI Gene Expression Omnibus (GEO) repository according to MIAME-compliant data submissions, NCBI GEO: GSE233916.

### Proteomic analysis

We carried out label-free quantification (LFQ) proteomics with sample preparation according to a published protocol with minor modifications ^83^. We used biological triplicates of Dataset 1: i) WT/WT iPSCs, ii) WT/WT NPCs, iii) 70Q/70Q iPSCs, iv) 70Q/70Q NPCs. We also used biological triplicates of Dataset 2: i) NGN2 neurons C1, ii) NGN2 neurons C2, iii) NGN2 neurons C3, iv) NGN2 neurons HD1, v) NGN2 neurons HD2, vi) NGN2 neurons HD3. We lysed all samples under denaturing conditions in buffer with 3 M guanidinium chloride (GdmCl), 5 mM tris-2-carboxyethyl-phosphine, 20 mM chloroacetamide, and 50 mM Tris-HCl at pH 8.5. We denatured lysates at 95°C for 10 min in a thermal shaker at 1000 rpm, and sonicated them in a water bath for 10 min. A small aliquot of cell lysate was used for the BCA assay to quantify protein concentration. We diluted the lysates (100 µg proteins) using a dilution buffer containing 10% acetonitrile and 25 mM Tris-HCl, pH 8.0, in order to reach a 1 M GdmCl concentration. We digested proteins with 1 µg LysC (MS-grade, Roche, Switzerland) while shaking at 700 rpm at 37°C for 2 h. The digestion mixture was diluted with the same dilution buffer in order to reach 0.5 M GdmCl. We added 1 µg trypsin (MS-grade, Roche, Switzerland) and incubated the digestion mixture in a thermal shaker at 700 rpm at 37°C overnight. We used solid phase extraction (SPE) disc cartridges (C18-SD, Waters, USA) for peptide desalting, according to the manufacturer’s instructions. We reconstituted desalted peptides in 0.1% formic acid in water and further separated them into four fractions by strong cation exchange chromatography (SCX, 3M Purification, Meriden, USA). We next dried the eluates in a SpeedVac, dissolved them in 20 µl 5% acetonitrile and 2% formic acid in water, briefly vortexed them, and sonicated them in a water bath for 30 sec prior to injection into nano-LC- MS. We carried out LC-MS/MS by nanoflow reverse phase liquid chromatography (Dionex Ultimate 3,000, Thermo Fisher Scientific, USA) coupled online to a Q-Exactive HF Orbitrap mass spectrometer (Thermo Fisher Scientific, USA). LC separation was performed with a PicoFrit analytical column (75 μm ID × 55 cm long, 15 µm Tip ID; New Objectives, Woburn, MA, USA) in-house packed with 3 µm C18 resin (Reprosil-AQ Pur, Dr. Maisch, Germany). We eluted peptides using a gradient from 3.8 to 40% solvent B in solvent A over 120 min at 266 nl per minute flow rate. Solvent A was: 0.1% formic acid, while solvent B was: 79.9% acetonitrile, 20% H2O, 0.1% formic acid. Nanoelectrospray was generated by applying 3.5 kV. A cycle of one full Fourier transformation scan mass spectrum (300−1750 m/z, resolution of 60,000 at m/z 200, AGC target 1e6) was followed by 12 data-dependent MS/MS scans (resolution of 30,000, AGC target 5e5) with a normalized collision energy of 25 eV. We used a dynamic exclusion window of 30 sec in order to avoid repeated sequencing of the same peptides. We sequenced only peptide charge states between two to eight. Raw MS data were processed with MaxQuant software (v1.6.10.43) and searched against the human proteome database UniProtKB with 20,600 entries, released in 05/2020. Parameters of MaxQuant database searching were: a false discovery rate (FDR) of 0.01 for proteins and peptides, a minimum peptide length of 7 amino acids, a mass tolerance of 4.5 ppm for precursor, and 20 ppm for fragment ions. We used the function “match between runs”. A maximum of two missed cleavages was allowed for the tryptic digest. We set cysteine carbamidomethylation as fixed modification, while N-terminal acetylation and methionine oxidation were set as variable modifications. We excluded any contaminants from further analysis, as well as any proteins identified by site modification or derived from the reversed part of the decoy database. We report the MaxQuant processed output files, peptide and protein identification, accession numbers, percentage sequence coverage of the protein, q-values, and LFQ intensities in **Supplementary Data 3, Data 4, Data 7**. We performed the correlation analysis of biological replicates and the calculation of significantly different metabolites and proteins using Perseus (v1.6.14.0). We transformed by log2 the LFQ intensities originating from at least two different peptides per protein group. We employed only protein groups with valid values within compared experiments. We carried out statistical analysis by a two-sample two-tailed t-test with Benjamini-Hochberg (BH, FDR of 0.05) correction for multiple testing. For comprehensive proteome data analyses, we applied gene set enrichment analysis (GSEA) ^84^. This was carried out to determine if a priori defined sets of proteins show statistically significant and concordant differences between mutations and controls. For GSEA analysis, we used all proteins with ratios calculated by Perseus. We applied GSEA standard settings, except that the minimum size exclusion was set to 5 and Reactome v7.2 and KEGG v7.2 were used as gene set databases. The cutoff for significantly regulated pathways was set to be ≤0.05 p-value and ≤0.25 FDR. We report the results of the GSEA analysis in **Supplementary Data 8**. The mass spectrometry proteomics data have been deposited to the ProteomeXchange Consortium via the PRIDE partner repository ^85^.

### Metabolomics analysis

Metabolite extraction and profiling by targeted LC-MS was performed as reported previously ^86^. We harvested biological triplicates of Dataset 2: i) NGN2 neurons C1, ii) NGN2 neurons C2, iii) NGN2 neurons C3, iv) NGN2 neurons HD1, v) NGN2 neurons HD2, vi) NGN2 neurons HD3. We aspirated the culture medium, quickly rinsed the cells twice with ice-chilled 1x PBS, pelleted the cells, and shock-froze them in liquid nitrogen. We extracted the metabolites with methyl tert-butyl-ether (MTBE), methanol, and water. The remaining protein pellets were used in BCA protein assay for normalization among samples. We aliquoted the extracts equally into additional tubes for later reconstitution in water, acetonitrile, and 50% methanol in acetonitrile. To each sample we added an internal standard mixture containing chloramphenicol, C13- labeled L-glutamine, L-arginine, L-proline, L-valine, and uracil (10 µM final concentration). We used a SpeedVac to dry the aliquots. We dissolved dry residuals in three different solvents: i) 100 µL 50% acetonitrile in MeOH with 0.1% formic acid, ii) 100 µL MeOH with 0.1% formic acid for analysis by HILIC column, or iii) 100 µL water, 0.1% formic acid for C18 column mode. We transferred the supernatants to micro-volume inserts. We injected 20 µL per run for subsequent LC-MS analysis. Over 400 metabolites were selected to cover most of the important metabolic pathways in mammals. Since metabolites are very diverse in their chemical properties, we used two different LC columns for metabolite separation: Reprosil-PUR C18-AQ (1.9 µm, 120 Å, 150 × 2 mm ID; Dr. Maisch, Germany) and zicHILIC (3.5 µm, 100 Å, 150 × 2.1 mm ID; Merck, Germany). We used the settings of the LC-MS instrument,1290 series UHPLC (Agilent Technologies, USA) online coupled to a QTrap 6500 (Sciex, USA) as reported previously ^87^. The buffer conditions were: A1, 10 mM ammonium acetate, pH 3.5 (adjusted with acetic acid); B1, 99.9% acetonitrile with 0.1% formic acid; A2, 10 mM ammonium acetate, pH 7.5 (adjusted with ammonia solution); B2, 99.9% methanol with 0.1% formic acid. We prepared all buffers in LC-MS grade water and organic solvents. We performed peak integration with MultiQuantTM software v.2.1.1 (Sciex, USA) and reviewed it manually. We normalized peak intensities, first against the internal standards, and subsequently against protein abundances obtained from the BCA assay. We used the first transition of each metabolite for relative quantification between samples and controls. Statistical analysis was carried out with Perseus ^88^. We employed a two-sample two-tailed t-test with Benjamini-Hochberg (BH, FDR of 0.05) correction for multiple testing. We provide the list of all metabolites including MRM ion ratios, KEGG and HMDB metabolite identifiers, and statistical values in **Supplementary Data 9**. These data were obtained using a previously reported LC-MS method containing the list of metabolites, transitions, and retention times ^87^. We deposited all original LC-MS generated QTrap wiff files on the peptide atlas repository with the identifier PASS04827, http://www.peptideatlas.org/PASS/PASS04827.

### Proteomic-driven functional metabolic analysis

For the functional metabolic analysis of the proteomic Dataset 1 (including iPSCs and NPCs from WT/WT and 70Q/70Q), we used the Quantitative System Metabolism (QSM) pipeline developed by Doppelganger Biosystem GmbH. We used the approach as described before ^35^. QSM data analysis used quantitative information on the expression levels of metabolic proteins (enzymes) to determine metabolic profiles, metabolic states and capacities, and metabolic fluxes. To ensure data quality, we used the quality control (QC) score, which counts the number of proteins of interest found, and the QSM score, which evaluates the number of metabolic processes associated with the enzymes found. For our dataset, the QC score was about 80% and the QSM score was around 100%, indicating excellent data quality that ensures reliable interpretability of the results. The kinetic model includes major cellular metabolic pathways of energy metabolism in neuronal cells, as well as key electrophysiological processes at the inner mitochondrial membrane, the mitochondrial membrane potential membrane, the transport of various ions, and the utilization of the proton motive force. Maximal enzyme activities (Vmax values) were estimated based on functional characteristics and metabolite concentrations of healthy neuronal tissues ^89^. Individual metabolic models were inferred using protein intensity profiles. The maximal activities (vmx) mean control for the normal state were previously calculated ^89^. QSM was used to calculate maximal energetic capacity. Energetic capacity was assessed under saturating glucose and oxygen concentrations, corresponds to healthy physiological conditions. Energetic capacities were evaluated by computing the changes of metabolic state elicited by an increase of the ATP consumption rate above the resting value. Maximal neuronal glucose uptake rate was assessed by increasing plasma glucose concentration in a systematic manner as the only energy delivering substrate, assuming high energy demands, and providing saturating oxygen concentrations.

### PCR analyses

For HTT PCR, genomic DNA was harvested from iPSCs colonies with Phire Animal Tissue Direct PCR Kit (Qiagen, USA). Exon 1 of the HTT locus was amplified by PCR using AmpliTaq Gold 360 DNA polymerase (Thermo Fisher Scientific, USA) and HTT specific primers using following PCR conditions: 95 °C 10 minutes, 35 cycles: 95 °C for 30 s, 55 °C for 30 s, and 72 °C for 1 min. Products were visualised by agarose gel electrophoresis, individual DNA bands were excised and purified using The Wizard® SV Gel and PCR Clean-Up System (Promega), cloned with CloneJET PCR Cloning Kit (Thermo Fisher Scientific, USA), and submitted to LGC Genomics for Sanger sequencing. Chromatograms were analysed using CLC Genomics Workbench (Qiagen, USA). For quantitative real-time PCR (qPCR) analysis of cerebral organoids, RNA extraction was performed from four cerebral organoids per condition with RNeasy kit (Qiagen, USA). cDNA was generated from 100 ng RNA using the first strand cDNA synthesis kit (Thermo Fisher Scientific, USA). qPCR experiments were conducted for three technical replicates and three independent biological replicates with the SYBR green Mastermix (Thermo Fisher Scientific). Amplification of 10 ng RNA was performed in a Viia7 Real Time PCR system (Thermo Fisher Scientific) as follows: 2 min activation step at 50°C, followed by 10 min at 95°C, 40 cycles of denaturation for 15 sec at 95°C, and annealing for 1 min at 60°C, finalized with 15 sec at 95°C, 1 min at 60°C, and 15 sec at 95°C. After averaging the CT-values of the three technical replicates, the expression levels of the genes of interest were normalized relative to the average expression of housekeeping genes GAPDH, OAZ1 and ACTB using the delta-delta-CT method. All primer sequences are reported in **Supplementary Data 10.**

### Immunostaining

NPCs and NGN2 neurons were grown on Matrigel-coated (Corning, USA) coverslips and fixed with 4% PFA (Science Services, Germany) for 20 min at RT and washed three times with PBS. For permeabilization, cells were incubated with blocking solution (PBS [Gibco, USA], 10% normal donkey serum, 1% Triton-X-100, 0.05% Tween-20 [all Sigma-Aldrich, USA]) for 1 h at RT. Primary antibodies were diluted in blocking solution and incubated overnight at 4°C on a shaker. Primary antibodies used were as follows: rabbit anti-CHCHD2 (1:800; Atlas Antibodies, Sweden) and mouse anti-TOM20 (1:500; MilliporeSigma, USA). Next, the cover slips were washed three times with PBS and incubated for 1 h at RT with Hoechst 33342 (1:2500; Life Technologies, USA) and the secondary antibodies in blocking solution at the following concentrations: donkey anti-rabbit Alexa Fluor 488 (1:300; Thermo Fisher Scientific, USA) and donkey anti-mouse Alexa Fluor 568 (1:300; Thermo Fisher Scientific, USA) Finally, cover slips were washed three times with PBS, mounted, air-dried and images were acquired using the confocal microscope Olympus Fluoview 3000 (Olympus, Japan).

Cerebral organoids were fixed with 4% PFA for 20 min at RT and washed 3x with PBS for 15 min. Next, organoids were cryoprotected overnight in a 30% sucrose in PBS solution. Following this, organoids were embedded in a 10% sucrose / 13 % gelatin solution in PBS and stored at -80°C upon further use. Hereafter, sections of 20 µm were cut using a Leica CM3050 S cryostat, collected on Superfrost Plus slides and stored at -80°C. Next, sections were washed three times with warm PBS for 10 sec to dissolve and get rid of any remaining gelatin. Subsequently, sections were fixed again with 4% PFA for 20 min at RT, washed 3x for 10 min with PBS, and blocked for 1 h at RT in blocking solution (PBS, 10% normal donkey serum, 1% Triton X-100). Following this, sections were incubated overnight at 4°C with the primary antibodies in blocking solution at the following concentrations: mouse anti-TUJ1 (1:2,000; Sigma-Aldrich, USA), goat anti-SOX2 (1:100; Santa Cruz Biotechnology, USA), rabbit anti- FOXA2 (1:300; Sigma-Aldrich, USA), goat anti-DARPP-32 (1:50; Santa Cruz Biotechnology, USA), rabbit anti-TOM20 (1:200; Santa Cruz Biotechnology, USA), guiney pig anti-GFAP (1:500; Synaptic Systems, Germany), mouse anti-SMI312 (1:500; BioLegend, USA), guiney pig anti-MAP2 (1:1,000; Synaptic Systems, Germany), mouse anti-EM48 (1:1,000; MilliporeSigma, USA), rabbit anti-Tau (1:500; Sigma-Aldrich, USA), mouse anti-Nestin (1:200; MilliporeSigma, USA), rabbit anti-PAX6 (1:200; BioLegend, USA), rabbit anti-CTIP2 (1:300; Abcam, UK), rabbit anti-FOXG1 (1:500; Abcam, UK) and rabbit anti-CHCHD2 (1:800, Atlas Antibodies, Sweden). Next, sections were washed 4x for 10 min with 0.1% Triton X-100 and 0.05% Tween-20 in PBS and incubated for 1-2 h at RT with Hoechst 33342 (1:2500; Life Technologies, USA) and the secondary antibodies in blocking solution at the following concentrations: donkey anti-mouse Alexa Fluor 488 (1:300; Thermo Fisher Scientific, USA), donkey anti-rabbit Alexa Fluor 488 (1:300, Thermo Fisher Scientific, USA), donkey anti-mouse Alexa Fluor 568 (1:300, Thermo Fisher Scientific, USA), donkey anti-rabbit CY3 (1:300; Jackson ImmunoResearch Laboratories, USA), donkey anti-rabbit CY5 (1:300; Jackson ImmunoResearch Laboratories, USA), donkey anti-goat CY5 (1:300; Jackson ImmunoResearch) and donkey anti-Guiney pig CY5 (1:300; Jackson ImmunoResearch Laboratories, USA). Finally, sections were washed 4x for 10 min with 0.1% Triton X-100 and 0.05% Tween-20 in PBS, mounted, air-dried and images were taken using the confocal microscopes Zeiss Z1 (Zeiss, Germany) and Olympus Fluoview 3000 (Olympus, Japan).

### CHCHD2 and TOM20 quantification

Images of NPCs and cerebral organoids stained against TOM20, CHCHD2, and Hoechst were processed as follows. We performed background subtraction using the rolling ball algorithm implementation of Scikit-image ^90^ with a radius of 50 pixels. To separate signal from background, we used Otsu’s thresholding method on the respective color channels. We then calculated the amount of colocalized pixels above the threshold for each marker. For comparing amounts of active TOM20 or CHCHD2 signals of different data sets, the amounts of positive signal per image were normalized by the amount of Hoechst signal per image to account for the fact that different contain different number of cells. Mann-Whitney U tests were used to test for differences between data sets (cell lines or treatments). All analysis code is available on GitHub (https://github.com/Scaramir/HD_colocalization). For mitochondrial network morphology assay of cerebral organoids, pictures of 100x magnification were taken of TOM20 stained sections using a Zeiss Z1 microscope. Next, the pictures were analyzed using ImageJ, as described before ^34^. Briefly, background was reduced, local contrast enhanced, pictures were made binary and using a “tubeness” plugin mitochondrial structures were tubed. After bandpass filtering, a threshold was set to minimize extremely small and large structures. Lastly, particles were analyzed, yielding information about the size and morphology of the structures.

### Western blotting and filter retardation assay (FRA)

Samples from NPCs and guided brain organoids were resuspended in 150 µL RIPA buffer (150 mM NaCl, 50 mM Tris, 0.5% w/v sodium desoxycholate, 1% v/v Triton X-100, 0.1% SDS, before using freshly added 1 mM PMSF and Protease Inhibitor Cocktail 1:100. Cell suspensions were pipetted 5x by using syringes to pass through 27G needles. The lysates were incubated at 4°C for 1 hr on a turning wheel followed by repeating the step of 27G needle treatment. The lysates were centrifuged at 15,000 x g for 15 min at 4°C. The supernatants were transferred to tubes and the protein concentrations were measured by using a BCA Protein Assay kit (Thermo Fisher Scientific, USA). For western blot, 5 µg proteins of each sample were used for detection. In these experiments, polyacylamide gels consisting of a 4% stacking gel and a 4-12% discontinuous resolving gel were applied. After SDS-PAGE, the proteins were transferred onto nitrocellulose membrane (0.45 µm) by tank blotting either at 500 mA, RT for 1.5 h or 100 mA, 4°C for 16 h. Proteins on the membranes were detected by using anti-HTT (Anti-Huntingtin antibody [EPR5526] (ab109115) Abcam; 1:5,000) or anti-GAPDH (1:1,000) primary antibodies followed by anti-rabbit-HRP or anti-mouse-HRP (1:10,000 for both antibodies) secondary antibodies, respectively. For Filter Retardation Assay (FRA), 3 µg protein of each sample was loaded. Both cellulose acetate membrane and nitrocellulose membrane were equilibrated in PBS/0.01% SDS. The dot blot apparatus was assembled with either membrane and the pump system was connected and turned on continuously. After washing the membranes with PBS, the samples were applied followed by two washing steps. The membranes were dried and proteins on the membranes detected with the same antibodies applied in western blot. The quantified figures were analyzed and all signals were evaluated in ImageJ. The HTT signals were compared to the signals of the internal control GAPDH. All experiments were repeated three times.

### Bioenergetic profiling

Live-cell assessment of cellular bioenergetics in NGN2 neurons was performed using Seahorse XF96 extracellular flux analyzer (Seahorse Bioscience, USA), as we described before ^71^. Briefly, 20,000 neurons were plated into each Matrigel-coated (Corning, USA) well of the XF96 well plates. Cells were maintained in the plates for 2 weeks. On the assay day, neurons were incubated at 37°C and 5% CO_2_ for 60 min to allow media temperature and pH to reach equilibrium before starting the simultaneous measurement of mitochondrial respiration (oxygen consumption rate, OCR) and anaerobic glycolysis (extracellular acidification rate, ECAR) using the sequential introduction of oligomycin, FCCP, and then rotenone plus antimycin A (all products at 1 µM; Sigma-Aldrich, USA). Normalization to DNA content in each well of the plate was performed using the CyQUANT Kit (Molecular Probes, USA). The supernatants were stored before and after the seahorse assay and used for lactate measurement using a Lactate Fluorometric Assay Kit (Biovision, USA).

### Statistical analysis

Data are expressed as mean and standard deviation (mean ± SD) where normality of the distribution could be verified, or as median and quartiles (median [1^st^;4^th^ quartiles]) otherwise. Significance was assessed using parametric tests (Student’s t-test, ANOVA) for normally-distributed data and non-parametric tests (Mann-Whitney U test, Kruskal-Wallis) when normal distribution could not be verified. Unless otherwise indicated, data were analyzed using GraphPad-Prism software (Prism 4.0, GraphPad Software, USA).

### Data availability

There are restrictions to the availability of the patient-derived iPSC lines due to nature of our ethical approval that does not support sharing to third parties and does not allow to perform genomic studies to respect the European privacy protection law. The datasets generated during this study are available, in cases data protection laws did not prevent the original datasets from being published:

