## Supplemental table legends for "Mutant Huntingtin impairs neurodevelopment in human brain organoids through CHCHD2-mediated neurometabolic failure"

### Supplementary Data

**Supplementary Data 1.** Differentially expressed genes (DEG) in iPSCs, NPCs, cerebral organoids (COs) at d28 and CO at d49 derived from WT/WT and 70Q/70Q (related to Fig. 3 and Supp Fig. 3)

**Supplementary Data 2.** Common genes expressed in iPSCs, NPCs, cerebral organoids (COs) at d28 and CO at d49 derived from WT/WT and 70Q/70Q (related to Fig. 3 and Supp Fig. 3)

**Supplementary Data 3.** Proteomics: differentially expressed proteins in iPSCs 70Q/70Q compared to iPSCs WT/WT (two-tailed t-test with Benjamini-Hochberg correction for multiple comparisons; FDR<0.05) (related to Figure 3).

**Supplementary Data 4.** Proteomics: differentially expressed proteins in NPCs 70Q/70Q compared to NPCs WT/WT (two-tailed t-test with Benjamini-Hochberg correction for multiple comparisons; FDR<0.05) (related to Figure 3).

**Supplementary Data 5.** Functional metabolic analysis from proteomics of iPSCs 70Q/70Q compared to iPSCs WT/WT and NPCs 70Q/70Q compared to NPCs WT/WT (related to Fig. 5).

**Supplementary Data 6.** Total RNA-sequencing: differentially expressed genes and pathways in NGN2 neurons from Ctrl iPSCs (C1, C2, C3) and HD iPSCs (HD1, HD2, HD3) (two-tailed Wald test with Benjamini-Hochberg correction for multiple comparisons; FDR<0.05) (related to Fig. 6).

**Supplementary Data 7.** Proteomics: differentially expressed proteins in NGN2 neurons from Ctrl iPSCs (C1, C2, C3) and HD iPSCs (HD1, HD2, HD3) (two-tailed Wald test with Benjamini-Hochberg correction for multiple comparisons; FDR<0.05) (related to Fig. 6).

**Supplementary Data 8.** Proteomics: gene set enrichment analysis (GSEA) in NGN2 neurons from Ctrl iPSCs (C1, C2, C3) and HD iPSCs (HD1, HD2, HD3) (two-tailed Wald test with Benjamini-Hochberg correction for multiple comparisons; FDR<0.05) (related to Fig. 6).

**Supplementary Data 9.** Metabolomics: differentially expressed metabolites in NGN2 neurons from Ctrl iPSCs (C1, C2, C3) and HD iPSCs (HD1, HD2, HD3) (two-tailed Wald test with Benjamini-Hochberg correction for multiple comparisons; FDR<0.05) (related to Fig. 6).

**Supplementary Data 10.** List of sequences and primers for PCR, QRT-PCR, siRNA knockdown, and CRISRP/eCas9 genome editing.
