## Supplementary material for "Mutant Huntingtin impairs neurodevelopment in human brain organoids through CHCHD2-mediated neurometabolic failure": Supp figures and legends

### Supplementary Data

**Supplementary Data 1.** Differentially expressed genes (DEG) in iPSCs, NPCs, cerebral organoids (COs) at d28 and CO at d49 derived from WT/WT and 70Q/70Q (related to Fig. 3 and Supp Fig. 3)

**Supplementary Data 2.** Common genes expressed in iPSCs, NPCs, cerebral organoids (COs) at d28 and CO at d49 derived from WT/WT and 70Q/70Q (related to Fig. 3 and Supp Fig. 3)

**Supplementary Data 3.** Proteomics: differentially expressed proteins in iPSCs 70Q/70Q compared to iPSCs WT/WT (two-tailed t-test with Benjamini-Hochberg correction for multiple comparisons; FDR<0.05) (related to Figure 3).

**Supplementary Data 4.** Proteomics: differentially expressed proteins in NPCs 70Q/70Q compared to NPCs WT/WT (two-tailed t-test with Benjamini-Hochberg correction for multiple comparisons; FDR<0.05) (related to Figure 3).

**Supplementary Data 5.** Functional metabolic analysis from proteomics of iPSCs 70Q/70Q compared to iPSCs WT/WT and NPCs 70Q/70Q compared to NPCs WT/WT (related to Fig. 5).

**Supplementary Data 6.** Total RNA-sequencing: differentially expressed genes and pathways in NGN2 neurons from Ctrl iPSCs (C1, C2, C3) and HD iPSCs (HD1, HD2, HD3) (two-tailed Wald test with Benjamini-Hochberg correction for multiple comparisons; FDR<0.05) (related to Fig. 6).

**Supplementary Data 7.** Proteomics: differentially expressed proteins in NGN2 neurons from Ctrl iPSCs (C1, C2, C3) and HD iPSCs (HD1, HD2, HD3) (two-tailed Wald test with Benjamini-Hochberg correction for multiple comparisons; FDR<0.05) (related to Fig. 6).

**Supplementary Data 8.** Proteomics: gene set enrichment analysis (GSEA) in NGN2 neurons from Ctrl iPSCs (C1, C2, C3) and HD iPSCs (HD1, HD2, HD3) (two-tailed Wald test with Benjamini-Hochberg correction for multiple comparisons; FDR<0.05) (related to Fig. 6).

**Supplementary Data 9.** Metabolomics: differentially expressed metabolites in NGN2 neurons from Ctrl iPSCs (C1, C2, C3) and HD iPSCs (HD1, HD2, HD3) (two-tailed Wald test with Benjamini-Hochberg correction for multiple comparisons; FDR<0.05) (related to Fig. 6).

**Supplementary Data 10.** List of sequences and primers for PCR, QRT-PCR, siRNA knockdown, and CRISRP/eCas9 genome editing.

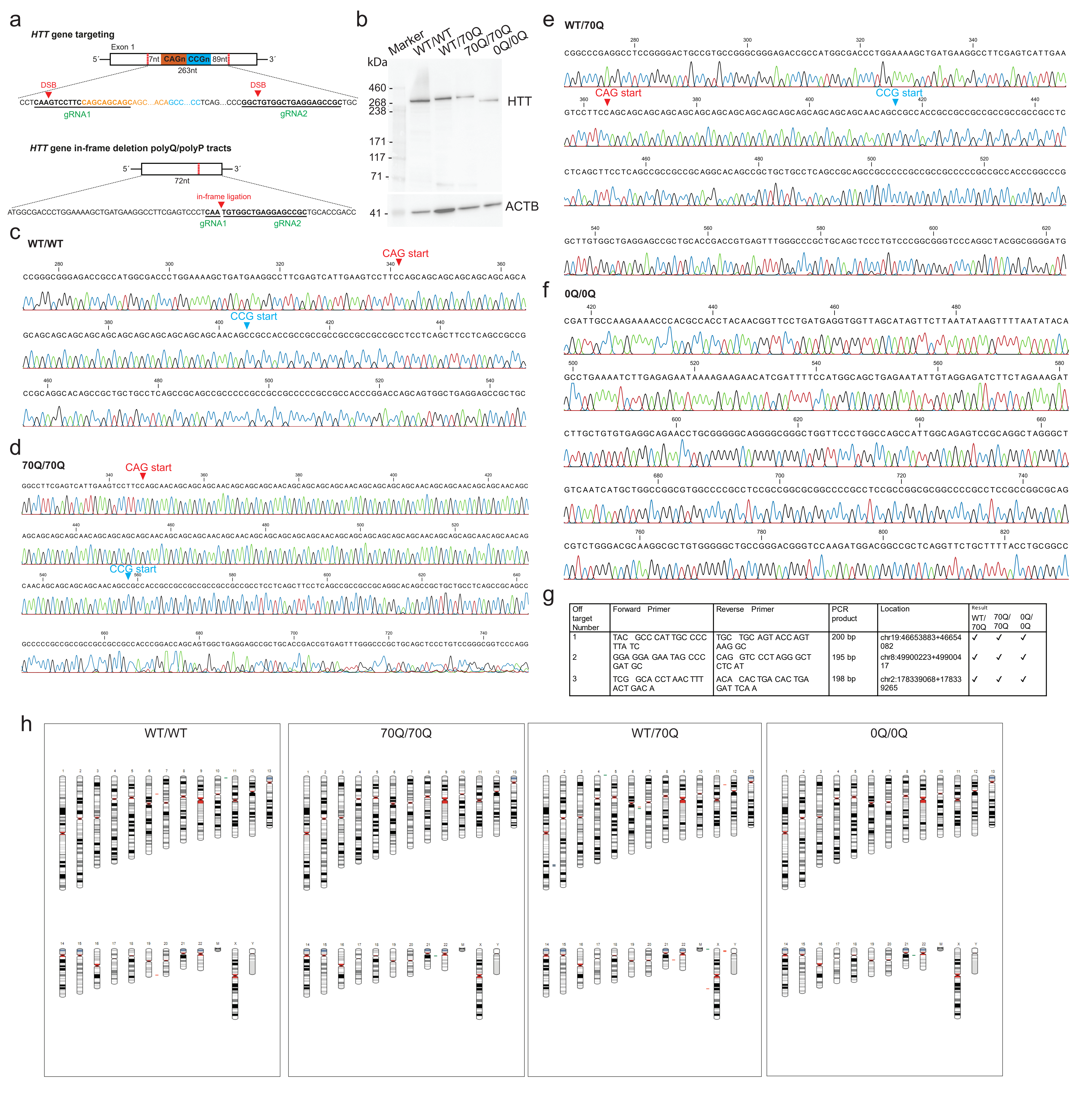


| Off target Number | Forward Primer | Reverse Primer | PCR product | Location | Result WT/70Q | 70Q/70Q | 0Q/0Q |
|---|---|---|---|---|---|---|---|
| 1 | TAC GCC CAT TGC CCC TTA TC | TGC TGC AGT ACC AGT AAG GC | 200 bp | chr19:46653883+46654082 | ✓ | ✓ | ✓ |
| 2 | GGA GGA GAA TAG CCC GAT GC | CAG GTC CCT AGG GCT CTC AT | 195 bp | chr8:49900223+49900417 | ✓ | ✓ | ✓ |
| 3 | TCG GCA CCT AAC TTT ACT GAC A | ACA CAC TGA CAC TGA GAT TCA A | 198 bp | chr2:178339068+178339265 | ✓ | ✓ | ✓ |

**Supplementary Fig. 1. Characterization of engineered isogenic iPSCs carrying mHTT (related to Fig. 1). a** Schematics of HTT gene targeting. DSB: double-strand breaks, gRNAs: guide RNAs. **b** Representative immunoblot image of engineered isogenic iPSC lines using anti-Huntingtin antibody (ab109115). **c-f** Sanger sequencing in the target HTT region (including indication of CAG and CCG starting sequences) showing one allele for WT/WT iPSCs and for each of the engineered isogenic iPSC lines (WT/70Q, 70Q/70Q, 0Q/0Q). **g** Off-target analysis of top three predicted sites by CRISPOR. Analysis in the engineered iPSC lines (WT/70Q, 70Q/70Q, 0Q/0Q) did not show any modification. **h** Normal karyotype assessed by SNP karyotyping (visualized with KaryoStudio v1.4) in WT/WT iPSCs and in the derived engineered lines (WT/70Q, 70Q/70Q, 0Q/0Q).

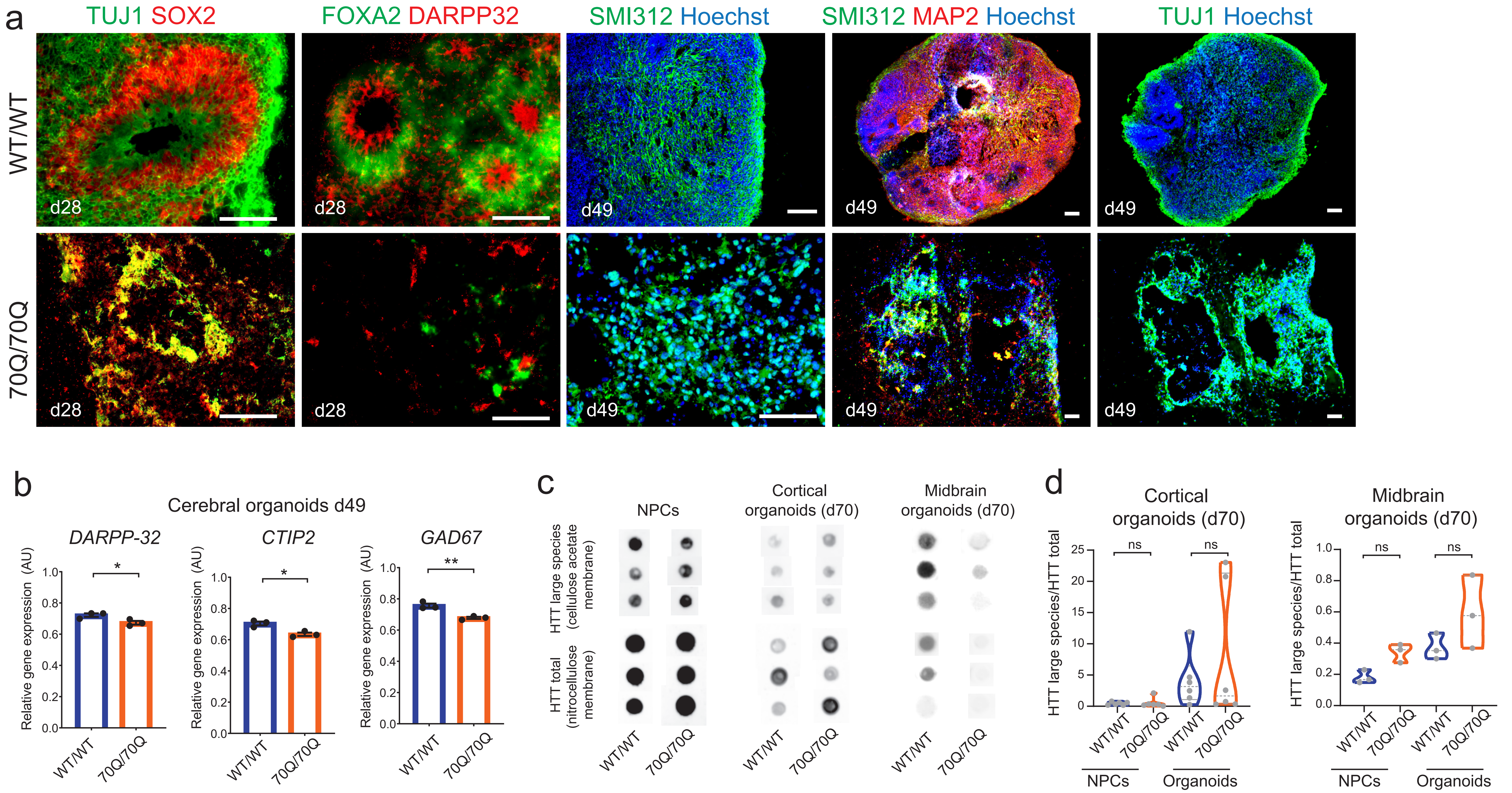

a
TUJ1 SOX2
FOXA2 DARPP32
SMI312 Hoechst
SMI312 MAP2 Hoechst
TUJ1 Hoechst
WT/WT
70Q/70Q
d28
d49
b
Cerebral organoids d49
DARPP-32
CTIP2
GAD67
Relative gene expression (AU)
WT/WT
70Q/70Q
c
NPCs
Cortical organoids (d70)
Midbrain organoids (d70)
HTT large species (cellulose acetate membrane)
HTT total (nitrocellulose membrane)
d
Cortical organoids (d70)
Midbrain organoids (d70)
HTT large species/HTT total
ns
NPCs
Organoids

**Supplementary Fig. 2. Characterization of brain organoids from engineered isogenic iPSCs carrying mHTT (related to Fig. 1-2). a** Immunostaining of unguided cerebral organoids at day 28 and 49 showing defective cytoarchitecture and neural progenitor cell (NPC) organization in 70/70Q. Scale bar: 100 µm. **b** qPCR analysis of neuronal markers in unguided cerebral organoids at day 49. Mean +/- s.e.m.; n=3 independent biological replicates (dots) per line; *$p<0.05$ **$p<0.01$; unpaired two-tailed t test. Four organoids were pooled for each individual RNA isolation. **c-d** Representative images and related quantification of filter retardation assay (FRA) in NPCs and in guided region-specific brain organoids (cortical and midbrain) to assess the potential presence of HTT protein aggregates using anti-Huntingtin antibody (ab109115). Cellulose acetate membranes were used to visualize aggregated HTT protein species, and nitrocellulose membrane to visualize total HTT protein species. The experiments were repeated three times.

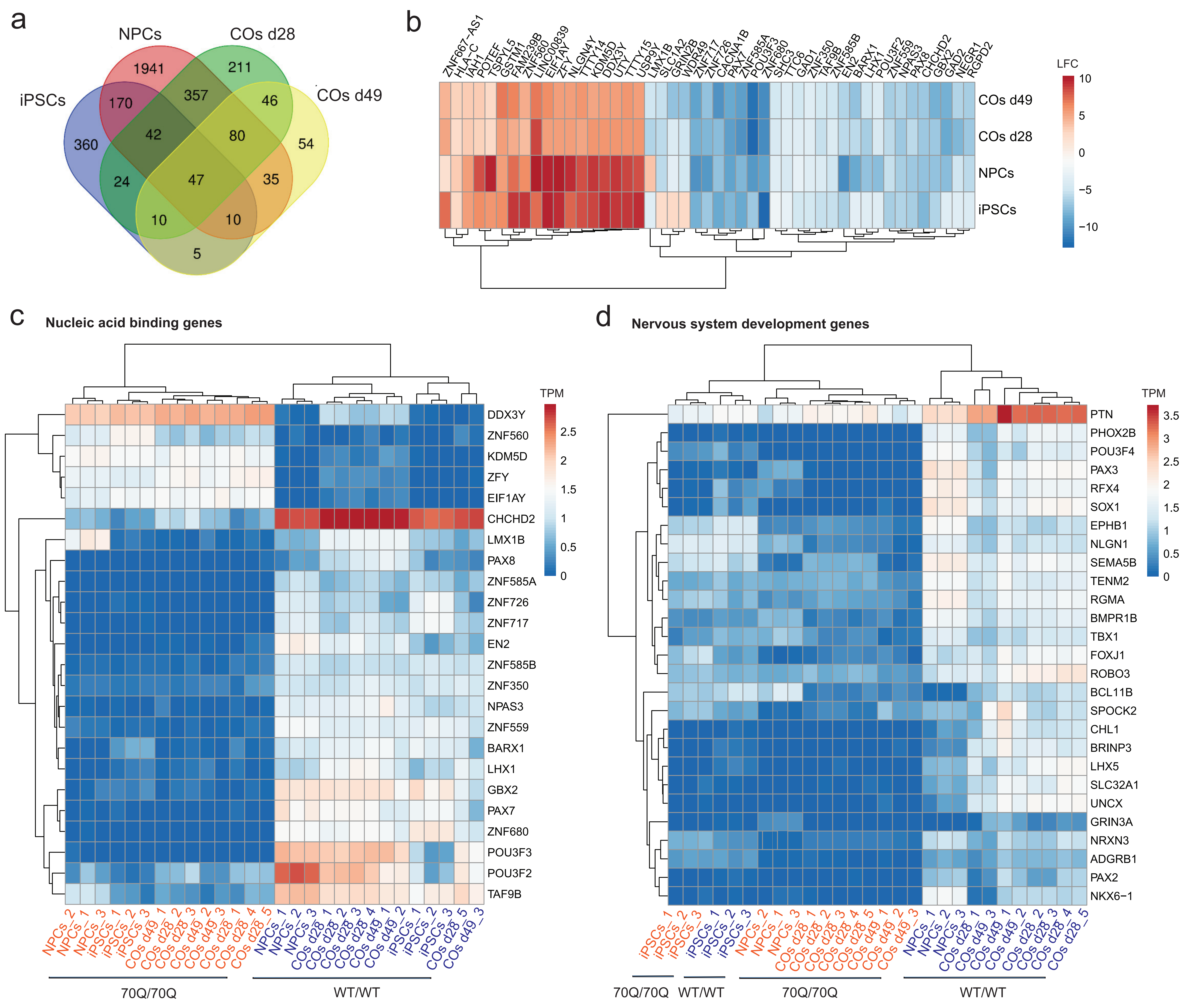
a
NPCs
COs d28
iPSCs
COs d49
1941
211
170
357
46
360
42
80
54
24
47
35
10
10
5
b
ZNF667−AS1
HLA−C
IAH1
POTEF
TSPYL5
GSTM1
FAM239B
ZNF560
LINC00839
EIF1AY
ZFY
NLGN4Y
TTTY14
KDM5D
DDX3Y
UTY
TTTY15
USP9Y
LMX1B
SLC1A2
GRIN2B
WDR49
ZNF717
ZNF726
CACNA1B
PAX7
ZNF585A
POU3F3
ZNF680
SHC3
TTC6
GAD1
ZNF350
TAF9B
ZNF585B
EN2
BARX1
LHX1
POU3F2
ZNF559
NPAS3
PAX8
CHCHD2
GBX2
GAD2
NEGR1
RGPD2
COs d49
COs d28
NPCs
iPSCs
LFC
10
5
0
−5
−10
c
Nucleic acid binding genes
TPM
2.5
2
1.5
1
0.5
0
DDX3Y
ZNF560
KDM5D
ZFY
EIF1AY
CHCHD2
LMX1B
PAX8
ZNF585A
ZNF726
ZNF717
EN2
ZNF585B
ZNF350
NPAS3
ZNF559
BARX1
LHX1
GBX2
PAX7
ZNF680
POU3F3
POU3F2
TAF9B
NPCs_2
NPCs_1
NPCs_3
iPSCs_1
iPSCs_2
iPSCs_3
COs d49_1
COs d28_2
COs d28_3
COs d49_2
COs d49_3
COs d28_1
COs d28_4
COs d28_5
NPCs_1
NPCs_2
NPCs_3
COs d28_1
COs d28_2
COs d28_3
COs d28_4
COs d49_1
COs d49_2
iPSCs_1
iPSCs_2
iPSCs_3
COs d28_5
COs d49_3
70Q/70Q
WT/WT
d
Nervous system development genes
TPM
3.5
3
2.5
2
1.5
1
0.5
0
PTN
PHOX2B
POU3F4
PAX3
RFX4
SOX1
EPHB1
NLGN1
SEMA5B
TENM2
RGMA
BMPR1B
TBX1
FOXJ1
ROBO3
BCL11B
SPOCK2
CHL1
BRINP3
LHX5
SLC32A1
UNCX
GRIN3A
NRXN3
ADGRB1
PAX2
NKX6−1
iPSCs_1
iPSCs_2
iPSCs_3
iPSCs_1
iPSCs_2
iPSCs_3
NPCs_2
NPCs_1
NPCs_3
COs d28_1
COs d28_2
COs d28_3
COs d28_4
COs d28_5
COs d49_1
COs d49_2
COs d49_3
NPCs_1
NPCs_2
NPCs_3
COs d28_1
COs d49_3
COs d49_1
COs d49_2
COs d28_2
COs d28_3
COs d28_4
COs d28_5
70Q/70Q
WT/WT
70Q/70Q
WT/WT

**Supplementary Fig. 3. Transcriptional signatures of engineered mHTT cells across different neurodevelopmental stages (related to Fig. 3). a** Venn diagram of unique and common transcripts among iPSCs, NPCs, cerebral organoids (COs) at day 28, and COs at day 49. **b** Heatmap of log-fold change (LFC) comparisons of differential gene expression of the 47 transcripts in common across all neurodevelopmental stages in 70/70Q cells compared to WT/WT cells. **c-d** Heatmap of the normalized expression values (TPM: Transcripts Per Million) of transcripts belonging to the GO category of “nucleic acid binding” and “nervous system development”.

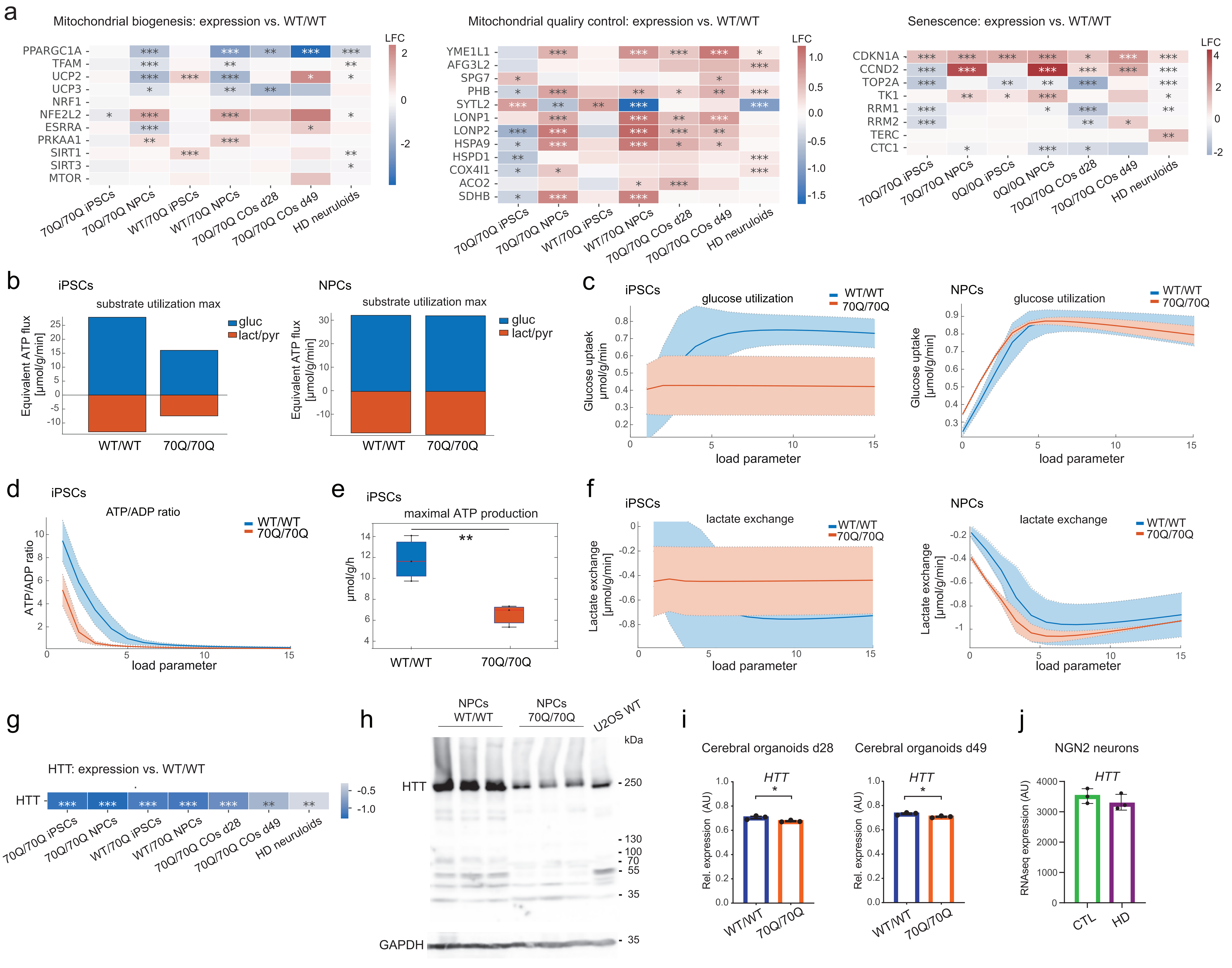
a
Mitochondrial biogenesis: expression vs. WT/WT
PPARGC1A
TFAM
UCP2
UCP3
NRF1
NFE2L2
ESRRA
PRKAA1
SIRT1
SIRT3
MTOR
LFC
70Q/70Q iPSCs
70Q/70Q NPCs
WT/70Q iPSCs
WT/70Q NPCs
70Q/70Q COs d28
70Q/70Q COs d49
HD neuruloids
Mitochondrial quality control: expression vs. WT/WT
YME1L1
AFG3L2
SPG7
PHB
SYTL2
LONP1
LONP2
HSPA9
HSPD1
COX4I1
ACO2
SDHB
Senescence: expression vs. WT/WT
CDKN1A
CCND2
TOP2A
TK1
RRM1
RRM2
TERC
CTC1
0Q/0Q iPSCs
0Q/0Q NPCs
b
iPSCs
substrate utilization max
Equivalent ATP flux [μmol/g/min]
gluc
lact/pyr
WT/WT
70Q/70Q
NPCs
c
iPSCs
glucose utilization
Glucose uptaek μmol/g/min
load parameter
NPCs
Glucose uptake [μmol/g/min]
d
iPSCs
ATP/ADP ratio
e
iPSCs
maximal ATP production
μmol/g/h
f
iPSCs
lactate exchange
Lactate exchange [μmol/g/min]
NPCs
g
HTT: expression vs. WT/WT
HTT
h
NPCs WT/WT
NPCs 70Q/70Q
U2OS WT
kDa
HTT
GAPDH
i
Cerebral organoids d28
Cerebral organoids d49
HTT
Rel. expression (AU)
j
NGN2 neurons
RNAseq expression (AU)
CTL
HD

**Supplementary Fig. 4. Metabolic defects in iPSCs and NPCs carrying mHTT (related to Fig. 4-5). a** Heatmap of log-fold change (LFC) comparisons of genes involved mitochondrial biogenesis, mitochondrial quality control, and senescence based on bulk RNA sequencing from iPSCs, NPCs, and cerebral organoids (COs) derived from 70Q/70Q and WT/70Q compared to WT/WT. HD neuruloids were compared to WT neuruloids. **b** Proteomics-driven functional metabolic analysis depicting the substrate utilization at maximal energy demands of iPSCs and NPCs from WT/WT and 70Q/70Q. Energetic capacities were evaluated by computing the changes of metabolic state elicited by an increase of the ATP consumption rate above the resting value. Bar graphs show consumption of glucose (gluc) and production of lactate-pyruvate (lact/pyr). **c** Computed glucose utilization rate in iPSCs and NPCs in dependence of increasing energetic demands. **d-e** Computed ATP/ADP ratio in dependence of increasing energetic demands and maximal ATP production capacity in iPSCs from WT/WT and 70Q/70Q. **f** Computed lactate exchange rate in iPSCs and NPCs in dependence of increasing energetic demands. **g** Heatmap of LFC comparisons of HTT gene based on bulk RNA sequencing performed in iPSCs, NPCs, and COs from 70Q/70Q and WT/70Q compared to WT/WT. HD neuruloids were compared to WT neuruloids. **h** Representative immunoblot of wild-type HTT in NPCs from WT/WT and 70Q/70Q. **i** qPCR-based HTT expression in unguided cerebral organoids at day 28 and 49. Mean +/- s.e.m.; n=3 independent biological replicates (dots) per line; *$p<0.05$ **$p<0.01$; unpaired two-tailed t test. Four organoids were pooled for each individual RNA isolation. **j** HTT expression based on RNA sequencing in NGN2 neurons derived from three controls (C1, C2, C3) and from three individuals with HD (HD1, HD2, HD3).

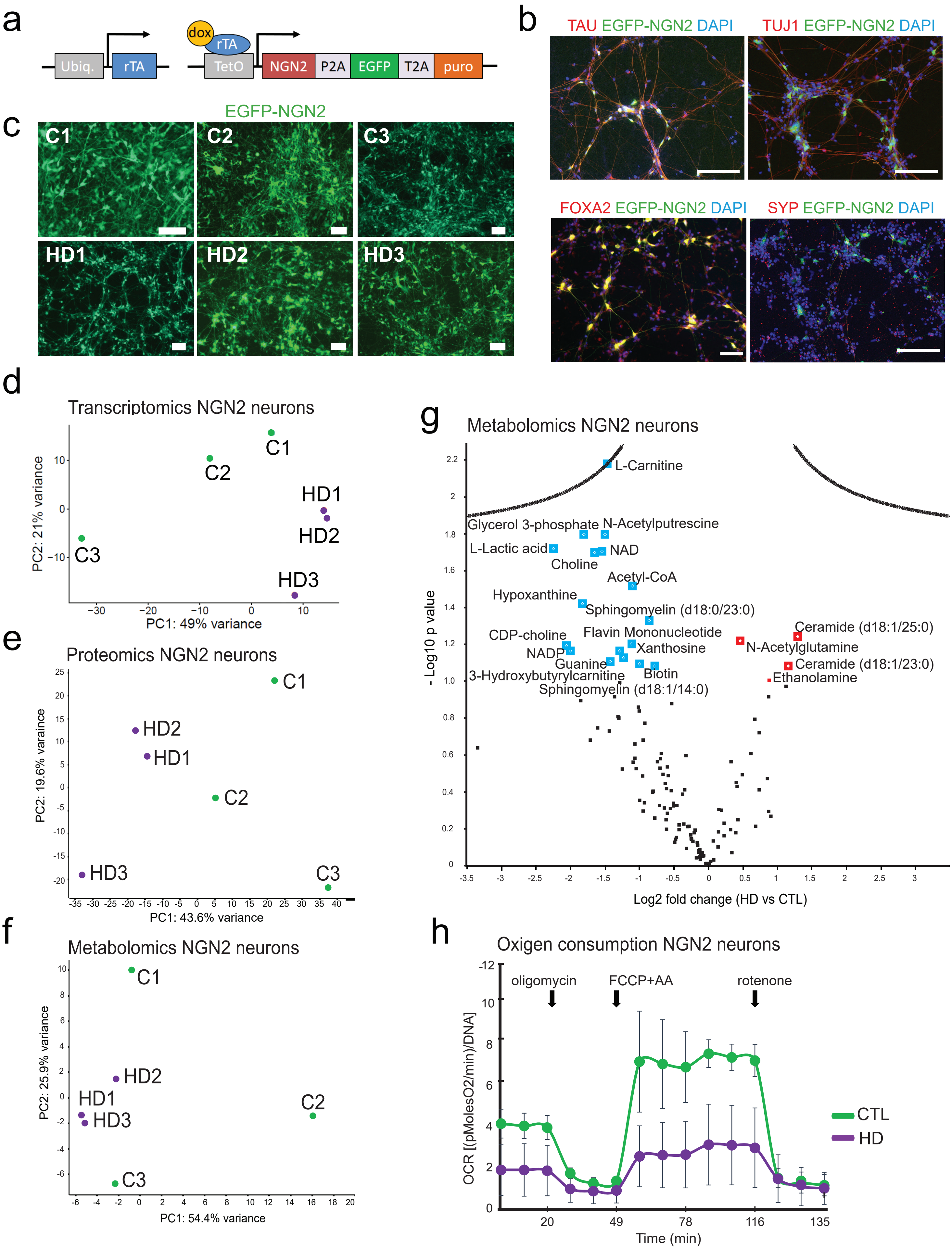
a
Ubiq.
rTA
dox
rTA
TetO
NGN2
P2A
EGFP
T2A
puro
b
TAU EGFP-NGN2 DAPI
TUJ1 EGFP-NGN2 DAPI
FOXA2 EGFP-NGN2 DAPI
SYP EGFP-NGN2 DAPI
c
EGFP-NGN2
C1
C2
C3
HD1
HD2
HD3
d
Transcriptomics NGN2 neurons
PC2: 21% variance
PC1: 49% variance
e
Proteomics NGN2 neurons
PC2: 19.6% varaince
PC1: 43.6% variance
f
Metabolomics NGN2 neurons
PC2: 25.9% variance
PC1: 54.4% variance
g
Metabolomics NGN2 neurons
- Log10 p value
Log2 fold change (HD vs CTL)
L-Carnitine
Glycerol 3-phosphate
N-Acetylputrescine
L-Lactic acid
NAD
Choline
Acetyl-CoA
Hypoxanthine
Sphingomyelin (d18:0/23:0)
CDP-choline
Flavin Mononucleotide
NADP
Xanthosine
Guanine
3-Hydroxybutyrylcarnitine
Biotin
Sphingomyelin (d18:1/14:0)
Ceramide (d18:1/25:0)
N-Acetylglutamine
Ceramide (d18:1/23:0)
Ethanolamine
h
Oxigen consumption NGN2 neurons
oligomycin
FCCP+AA
rotenone
OCR [(pMolesO2/min)/DNA]
Time (min)
CTL
HD

**Supplementary Fig. 5. Charachterization of NGN2 neurons from healthy controls and individuals with HD (related to Fig. 6). a** Lentiviral constructs used for transducing NPCs. **b** Expression of neuronal and synaptic markers in NGN2 neurons at day 14 of differentiation from NPCs. Scale bar: 100 µm. **c** NGN2 neurons at day 14 of differentiation from healthy controls (C1, C2, C3) and from individuals with HD (HD1, HD2, HD3). **d-f** Principal component analysis (PCA) plots showing the distribution of the NGN2 neurons from healthy controls (C1, C2, C3) and from individuals with HD (HD1, HD2, HD3) based on RNA sequencing, global proteomics, and metabolomics. **g** Metabolomics of NGN2 neurons. Key downregulated metabolites in HD neurons are highlighted in blue, key upregulated metabolites are highlighted in red. **h** Oxygen consumption rate (OCR) profile obtained with Seahorse Bioanalyzer in NGN2 neurons from healthy controls (C1, C2, C3) and from individuals with HD (HD1, HD2, HD3). Mean +/- s.e.m.; n=3 independent biological replicates per line.
